# ASNA-1 oxidation induced by cisplatin exposure enhances its cytotoxicity by selectively perturbing tail anchored protein targeting

**DOI:** 10.1101/821728

**Authors:** Dorota Raj, Ola Billing, Agnieszka Podraza, Oskar Hemmingsson, Gautam Kao, Peter Naredi

**Author notes:** These authors contributed equally (Dorota Raj, Ola Billing, Agnieszka Podraza).

## Abstract

Cisplatin is a frontline cancer treatment, but intrinsic or acquired resistance is common. We previously showed that ASNA-1/TRC40 inactivation increases cisplatin sensitivity in mammalian cells and a *Caenorhabditis elegans asna-1* knockdown model. ASNA-1 has conserved tail-anchored protein (TAP) targeting and insulin secretion functions. Here we examined the mechanism of ASNA-1 action. We show that ASNA-1 exists in two physiologically-responsive redox states with separable TAP-targeting and insulin secretion functions. Cisplatin-generated ROS targeted ASNA-1 oxidation, resulting in a selective targeting defect of an ASNA-1-dependent TAP. Increased ASNA-1 oxidation sensitized worms to cisplatin cytotoxicity. Mutants with a redox balance favoring oxidized ASNA-1 were cisplatin sensitive as null mutants by diverting ASNA-1 away from its TAP-targeting role and instead perturbing endoplasmic reticulum (ER) function. Mutations in the ASNA-1 receptor required for TAP insertion induced equivalent cisplatin sensitivity. We reveal a previously undescribed cellular dysfunction induced by cisplatin, identify a cisplatin target, and show that drug exposure causes TAP targeting-induced ER dysfunction. Therapeutic oxidation of ASNA-1 could be a clinically useful means to increase cisplatin sensitivity, reduce cytotoxic drug doses, and counteract cisplatin resistance.

**AUTHOR SUMMARY:** Cisplatin is a very effective anti-cancer drug and is widely used as a frontline treatment. However, tumor resistance limits its use. Tumor re-sensitization would improve cancer treatment. ASNA-1/TRC40 knockdown in *Caenorhabditis elegans* and mammals results in cisplatin hypersensitivity, but the underlying mechanistic details are largely unknown. We show that in *C. elegans* ASNA-1 mutants, increased cisplatin killing is coupled with delocalization of a tail-anchored protein, SEC-61β, a membrane protein that should reach the ER and is instead mistargeted. Like its homologs, the reduced form of worm ASNA-1 is needed for targeting activity. Targeting is blocked upon ASNA-1 oxidation after cisplatin treatment, likely via reactive oxygen species (ROS) generated by cisplatin treatment. Nevertheless, the oxidized form of the protein can execute other functions like insulin secretion. We show also that mutants with high oxidized ASNA-1 levels are cisplatin sensitive. Additionally, cisplatin induced mistargeting strictly acts through ASNA-1 inactivation. Thus, we define a pathway from cisplatin exposure that targets protein (ASNA-1) inactivation, consequently leading to mis-targeting of proteins that need ASNA-1 for their maturation. This multi-step process provides vital information about likely proteins that can be targeted by drugs to enhance cisplatin mediated killing and improve chemotherapy.

**SUMMARY STATEMENT:** Sensitizing tumors to cisplatin would be of considerable therapeutic benefit. Here we show a novel mechanism of cisplatin sensitization via oxidation of ASNA-1 in a *Caenorhabditis elegans* model.

## INTRODUCTION

Solid tumors consist of both dividing and non-dividing post-mitotic cells (Komlodi-Pasztor et al., 2012), and both populations must be eliminated to reduce tumor bulk. The frontline cytotoxic chemotherapy cisplatin rapidly kills both dividing and non-dividing cells but via different mechanisms (Stewart, 2007). In dividing cells, cisplatin directly binds to DNA to cause DNA damage, cell cycle arrest, and apoptosis (Kelland, 2007; Legin et al., 2014). In non-dividing cells, it kills by binding to antioxidants like glutathione, elevating reactive oxygen species (ROS) levels that have downstream effects on a variety of cellular processes (Dasari and Tchounwou, 2014). Identification of proteins that are rapidly oxidized after cisplatin exposure would provide important information on the mechanism of cisplatin-induced death in post-mitotic cells. If cytotoxicity requires the oxidation of specific proteins, their targeted oxidation could be combined with lower doses of cisplatin to achieve equivalent anti-tumor activity as high-dose single agent regimens. The longstanding limitation of cisplatin use is the eventual development of tumor resistance and significant side-effects of nephrotoxicity and ototoxicity. The use of lower but effective doses would address these important limitations.

ASNA-1/TRC40/GET3 is a redox-sensitive protein whose knockdown increases the sensitivity of tumor cells to cisplatin-induced death (Hemmingsson et al., 2009). We previously established *Caenorhabditis elegans* as a relevant system to model the cisplatin killing effect in mammalian post-mitotic cells, since *C. elegans asna-1* mutants are sensitive to cisplatin-induced death (Hemmingsson et al., 2010). Human ASNA-1/TRC40 can act as a functional substitute in worm mutants, further underlining the model’s similarity to human cells (Kao et al., 2007). We have also shown that ASNA-1 in mouse and *C. elegans* promotes insulin secretion and that targeted mouse knockouts develop type 2 diabetes (T2D) (Kao et al., 2007; Norlin et al., 2016). This work led us to propose that the cisplatin response and insulin secretion functions are separable (Hemmingsson et al., 2010).

A possible molecular basis for the separation of functions emerges from extensive cellular and structural biology studies on ASNA-1 homologs in yeast, mammals, and plants (Mateja et al., 2009; Mateja et al., 2015; Srivastava et al., 2016; Voth et al., 2014). The yeast homolog GET3 can adopt two alternative redox-sensitive forms with non-overlapping functions (Voth et al., 2014). The best understood function is of the reduced dimeric form, which acts as part of a targeting complex to guide tail-anchored membrane proteins (TAPs) for insertion into the endoplasmic reticulum (ER) membrane (Favaloro et al., 2008; Schuldiner et al., 2008; Stefanovic and Hegde, 2007). However, GET3 can also act as a general chaperone and a holdase by adopting an oxidized tetrameric structure with internal disulfide bonds via oxidation of critical redox-sensitive cysteines (Powis et al., 2013; Voth et al., 2014). *In vitro*, GET3 adopts a zinc-stabilized conformation needed for TAP targeting when conserved cysteines are reduced but acts as a holdase and general chaperone when these residues are oxidized (Voth et al., 2014). Thus, a better understanding of how ASNA-1 works and which form of the protein should be targeted in different pathologies requires matching the disease with the molecular state before rational drug regimens can be devised.

Here we investigated links between cisplatin-induced oxidative stress, the redox-responsive nature of ASNA-1, and the role of ASNA-1 in cisplatin cytotoxicity. ASNA-1 is needed for ER targeting of a model TAP via its ER-localized receptor WRB-1. Specifically, we asked whether ASNA-1 activity in worms can be targeted and modified by cisplatin exposure and whether this would explain its role as a sensitivity factor. We show that reduced monomeric ASNA-1 (ASNA-1^RED^) protects the organism from cisplatin cytotoxicity. Cisplatin generates ROS, and drug exposure rapidly leads to higher levels of oxidized ASNA-1 (ASNA-1^OX^) at the expense of ASNA-1^RED^ and cisplatin treatment in wild-type worms perturbs model TAP targeting in an ASNA-1 dependent manner. Cisplatin-induced oxidation of ASNA-1 is not a bystander effect. Rather, mutants with increased levels of oxidized ASNA-1 are sensitive to cisplatin death. We find that increased oxidation of ASNA-1 is equivalent to complete depletion of ASNA-1 with respect to cisplatin response without having any effect on its insulin secretion function. Increased levels of oxidized ASNA-1 also re-distributes the protein away from its juxta-membrane localization. Failure of TAP targeting is important for cisplatin-induced death, because *wrb-1^−/−^* mutants are also cisplatin sensitive. Thus, we delineate a pathway in which cisplatin-induced ROS generation inactivates the cisplatin response function of ASNA-1, which in turn perturbs the targeting of a TAP protein to the ER membrane. Taken together, our studies identify ASNA-1 as a biologically relevant cisplatin target, reveal which molecular form of ASNA-1 is modulated, and demonstrates that one strategy to increase cisplatin sensitivity would be biasing ASNA-1 to the oxidized state.

## RESULTS

### ASNA-1 is present in physiologically redox-sensitive states

The oxidation state of the *S. cerevisiae* ASNA-1 homolog, GET3, modulates its activities. We therefore asked whether *C. elegans* ASNA-1 could also act as a redox-regulated switch and whether this switch responds to internal physiological changes. We performed non-reducing SDS-PAGE on proteins isolated from wild-type worms (**Fig. 1A**) and control worms expressing ASNA-1::GFP (**Fig. 1B-H**), and found that *C. elegans* ASNA-1 was present in both its oxidized and reduced states (**Fig. 1A-H**). GFP alone was not detected in the oxidized state (**Fig. 1H**). Consistent with the findings in yeast (Voth et al., 2014), there was no oxidation of ASNA-1::GFP when two conserved cysteines at positions 285 and 288 were substituted for serines (**Fig. 1G**). Moreover, the balance between ASNA-1:GFP^OX^ and ASNA-1:GFP^RED^ could be altered by exposure of worms to different pro-oxidants: 10 mM H_2_O_2_/1mM CuSO_4_ (**Fig. 1B**) or 5 mM sodium arsenite (**Fig. 1C**). ASNA-1:GFP^OX^ was diminished by exposing animals to the antioxidant mitoTempo (**Fig. 1D**). The balance shifted towards more ASNA-1:GFP^OX^ at the expense of ASNA-1:GFP^RED^ in *sod-2(gk257)* and *mev-1(kn1)* mutants, which have high endogenous ROS levels (**Fig. 1E,F**) (Labuschagne et al., 2013). We concluded that ASNA-1 exists in two alternative molecular states and that their balance can be altered by environmental factors and internal physiological changes.

**Figure 1.**
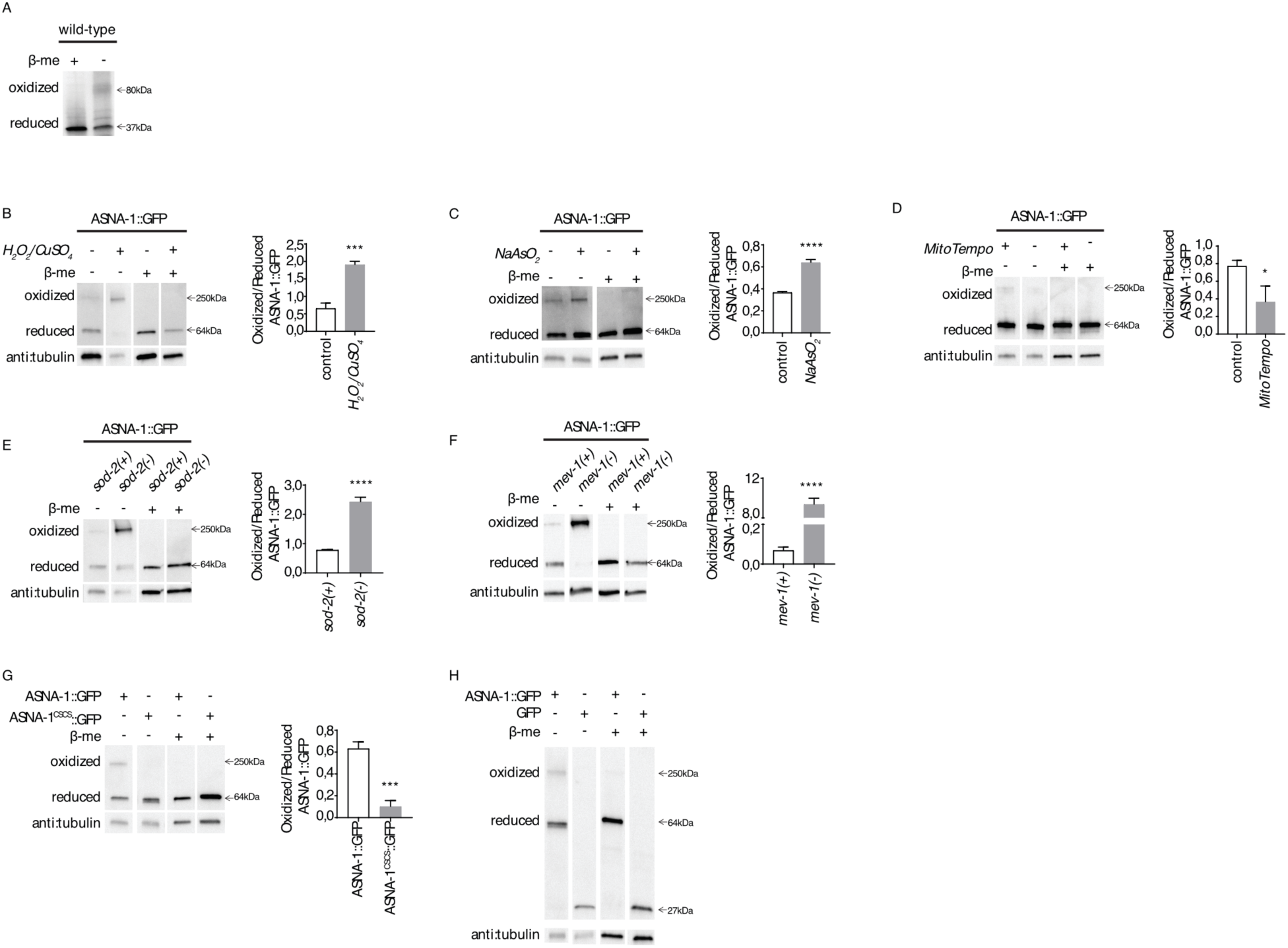
ASNA-1 is present in oxidized and reduced states. (A) Reducing and non-reducing SDS-PAGE visualizes oxidized and reduced ASNA-1 in the N2 strain. The blot was probed with anti-ASNA-1 antibody. Reducing and non-reducing SDS-PAGE visualizes oxidized and reduced ASNA-1::GFP: (B) exposed to 10 mM H_2_O_2_/1 mM CuSO_4_ for 30 min; (C) exposed to 5 mM NaAsO_2_ for 1 h; (D) exposed to the antioxidant MitoTempo 50 µM for 48 h; (E) in *sod-2(gk257)* mutants; (F) in *mev-1(kn1)* mutants; (G) in ASNA-1^C285S;C288S^::GFP transgenics; (H) in GFP-expressing strain *(unc-119(ed3);oxTi880*). Blots were probed with anti-GFP antibody and tubulin was used as a loading control. Every panel represents a single gel. Statistical significance was determined by the independent two-sample t-test (n ≥ 3; *p < 0.05, **p < 0.01, ***p < 0.001). Bars represent mean ± SD.

### ASNA-1 promotion of TAP insertion is independent of its role in insulin secretion

We next examined whether redox-induced structural changes in ASNA-1 impacted its well-studied functions and whether these functions were affected by alterations of the cellular ASNA-1 redox balance. While the ASNA-1/TRC40 pathway guides TAPs to the ER membrane, other pathways (EMC, HSP70/HSC40, SND) can potentially compensate for the absence of ASNA-1 to produce little overall defect in TAP targeting (Aviram and Schuldiner, 2017). To investigate whether the TAP targeting function was affected by the absence of worm ASNA-1, we established an *in vivo* model for TAP targeting in *C. elegans.* In vertebrates and yeast, SEC61β is a model TAP that requires ASNA1/TRC40 for correct targeting to the ER membrane (Favaloro et al., 2008; Hegde and Keenan, 2011; Schuldiner et al., 2008; Stefanovic and Hegde, 2007) via binding of ASNA-1 homologs to the transmembrane domain (TMD) of SEC61β. This domain is highly conserved in the *C. elegans* homolog *sec-61.B* (hereafter called SEC61β). Indeed, Co-IP/MS/MS analysis detected SEC61β as an ASNA-1::GFP-interacting partner (**Table 1**). Cell fractionation using GFP-tagged SEC61β confirmed that the protein was found only in the membrane fraction, even when expressed under a strong intestine-specific promoter (**Fig. 2F**). Co-expression of GFP::SEC-61β in intestinal cells of wild-type *C. elegans* with the mCherry-tagged rough ER (RER)-specific protein, SP12 (*spcs-1*), resulted in near complete colocalization in a pattern characteristic of ER (**Fig. 2A,B** and **Fig. S1**). This colocalization pattern was disrupted in animals expressing GFP::SEC61β^ΔTMD^ with its TMD deleted. Instead, large aggregates of GFP::SEC61β^ΔTMD^ accumulated in cytoplasmic regions distinct from the RER (**Fig. 2A,B**). Taken together, this established that worm SEC-61β was an ASNA-1-interacting partner that localized to the ER membranes via its TMD and thus was a good model protein to further study the contribution of ASNA-1 to TAP targeting. This localization required ASNA-1 since, in the two worm *asna-1* protein null mutants *(ok938* and *sv42)* (Kao et al., 2007), the ER localization of GFP::SEC-61β was significantly decreased and the protein was instead detected in cytoplasmic foci (**Fig. 2A,B**). The targeting defect was solely due to lack of ASNA-1, since wild-type ASNA-1 expressed from a transgene completely rescued the TAP targeting defect (**Fig. 2A,B** and **Fig. S1**). Detection of the same phenotype in two independent deletion mutants and rescue with wild-type ASNA-1 ruled out any contribution from background genetic effects.

**Figure 2.**
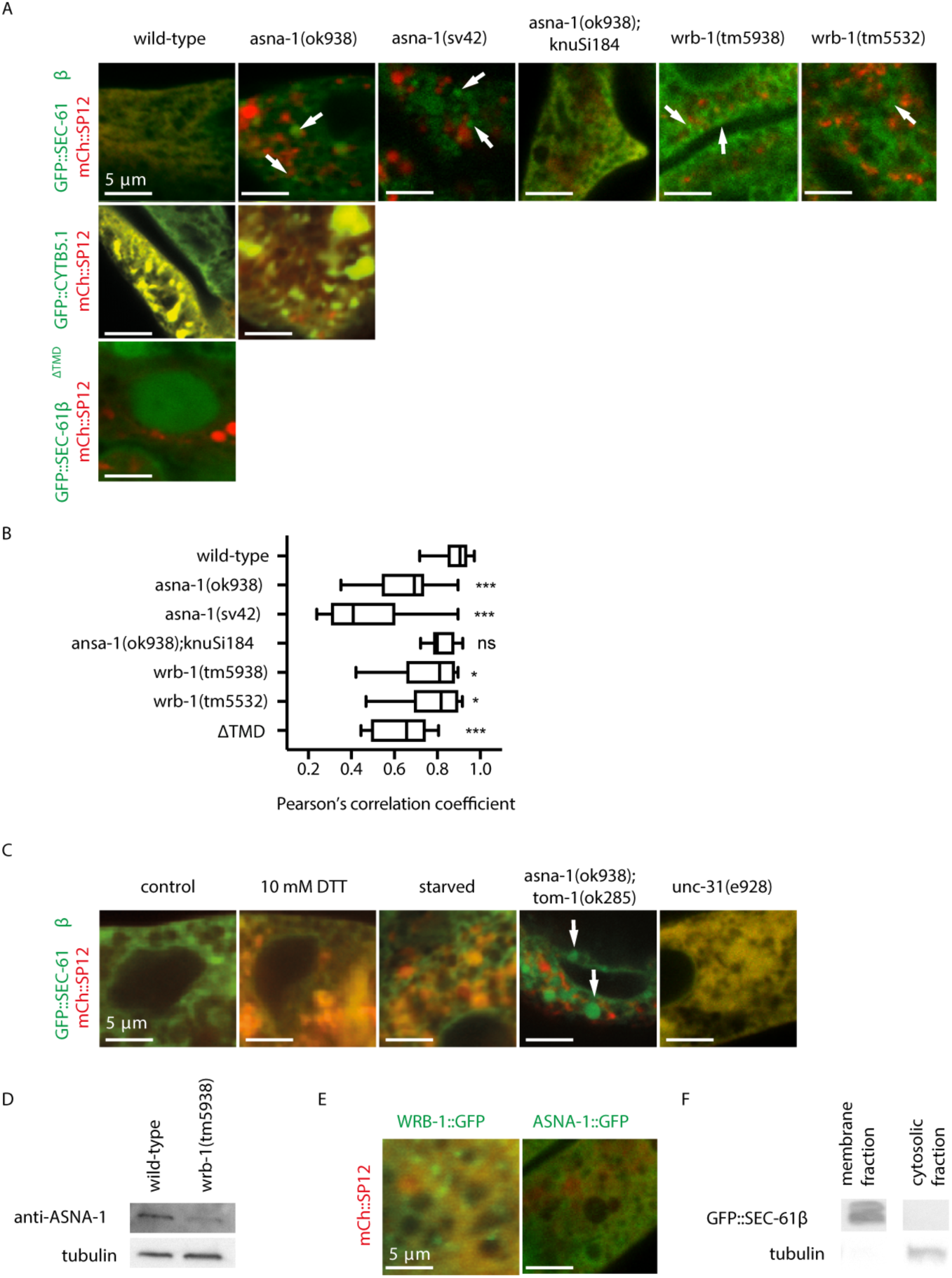
ASNA-1 and WRB-1 promote ER targeting of TAPs *in vivo*. (A) Confocal imaging merge of wild-type, *asna-1(ok938), asna-1(sv42), asna-1(ok938);knuSi184, wrb-1(tm5938)*, and *wrb-1(tm5532)* animals co-expressing GFP::SEC-61β or GFP::CYTB5.1 or GFP::SEC-61β^ΔTMT^ with mCherry::SP12. White arrows indicate GFP::SEC-61β foci that do not co-localize with mCherry::SP12. (B) Pearson’s correlation analysis of GFP::SEC-61β and mCherry::SP12 co-localization in different strains. Box plot represents the average Pearson correlation coefficient (R) of the indicated strains. Statistical significance was determined by one-way ANOVA followed by Bonferroni post-hoc correction (n ≥ 10; *p < 0.05, **p < 0.01, ***p < 0.001). (C) Confocal imaging merge of animals co-expressing GFP::SEC-61β and mCherry::SP12 with the indicated treatment and genetic backgrounds. White arrows label GFP::SEC-61β-positive foci that do not co-localize with mCherry::SP12. (D) Western blot analysis of ASNA-1 levels in wild-type and *wrb-1(tm5938)* animals. Blot was probed with anti-ASNA-1 antibody and tubulin was used as a loading control. (E) Confocal imaging merge of animals co-expressing WRB-1::GFP or ASNA-1::GFP and mCherry::SP12. (F) Subcellular localization of GFP::SEC-61β. Blot was probed with anti-GFP antibody. Tubulin used as a loading control.

**Table 1.**
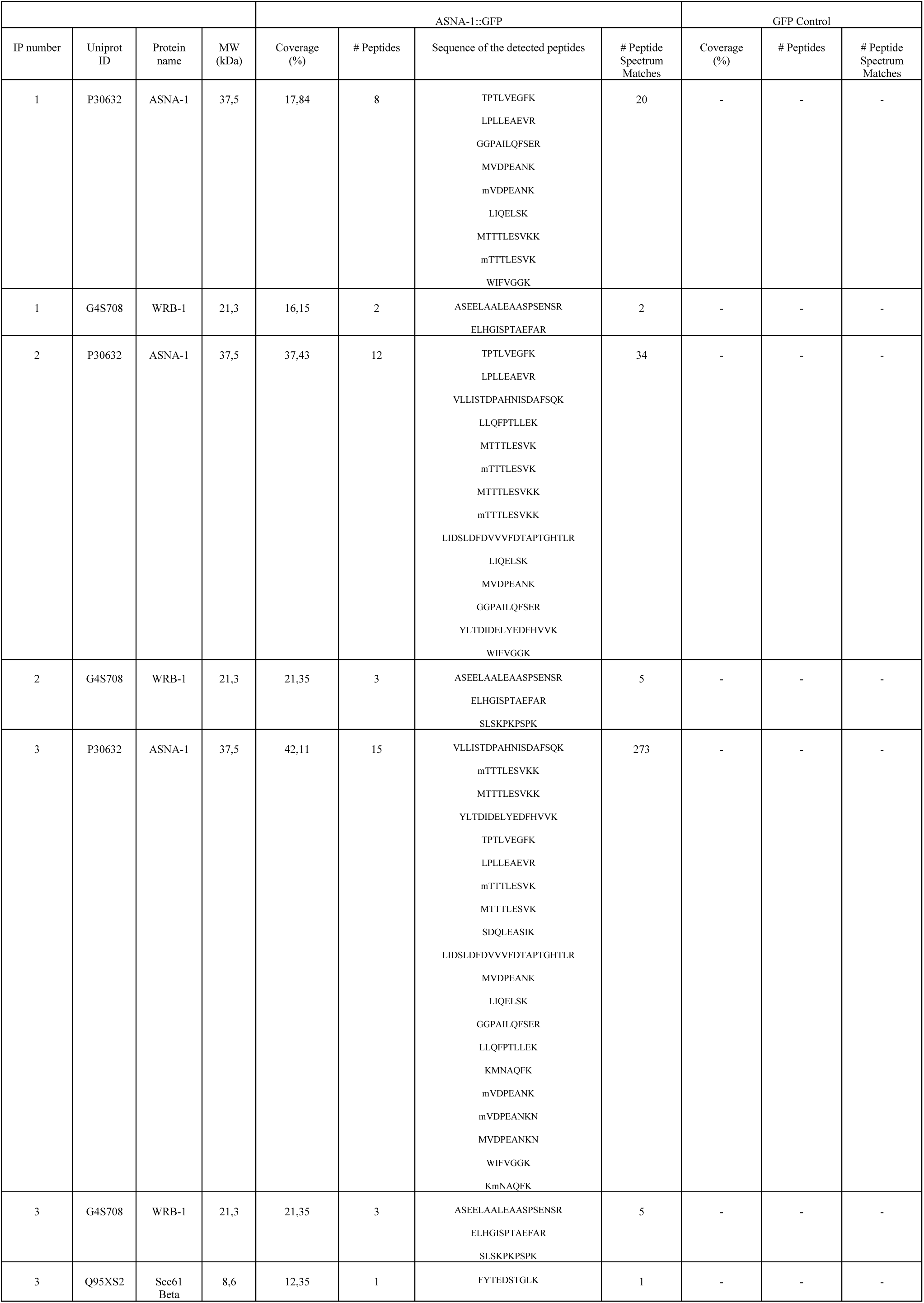
Proteomic identification of ASNA-1 interacting partners. Proteomic analysis of ASNA-1::GFP and GFP control pull down. Table shows: immunoprecipitation number (IP number), accession number of detected protein from UNIPROT (Uniprot ID), protein name detected in the analysis (Protein name), molecular weight of a detected protein (MW (kDa)), percentage of the protein sequence covered by the identified peptides (Coverage (%)), total number of peptides identified in the protein/protein group (# Peptides), sequences of the detected peptides, number of peptide spectrum matches for the peptide (# Peptide Spectrum Matches).

TEM analysis of *asna-1(ok938)* mutants revealed aggregates containing misfolded proteins and dilated RER lumina (**Fig. S2A,S2B**). The ER membrane was also found in autophagosomes (**Fig. S2C**), an ER stress-related structure, very similar to that seen in ASNA-1-deficient mice (Norlin et al., 2016). As expected, a transgene-based assay also revealed that autophagy levels were high in asna*-1(ok938)* mutants (**Fig. S2D**). The abnormal RER membranes in mutant animals were competent for proper membrane targeting of another TAP, the worm homologue of CytB5 (*cytb-5.1)*, an ASNA-1-independent, ER-specific TAP (Abell et al., 2007; Stefanovic and Hegde, 2007), which localized to the ER in *asna-1(ok938)* mutants to the same extent as in wild-type animals (**Fig. 2A, 6B**). This was also true for another ASNA-1-independent TAP, SERP-1.1(F59F4.2) (**Fig. 6A,B**). These findings further underscored the validity of our *in vivo* model, since it enabled us to distinguish between ASNA-1-dependent and independent TAP insertion.

*asna-1* mutants have multiple phenotypes including elevated autophagy levels (**Fig. S2D**), high ER stress (Natarajan et al., 2013), and low insulin signaling levels (Hemmingsson et al., 2010; Kao et al., 2007). We asked whether these phenotypes contribute to the TAP targeting defect in *asna-1(ok938)* mutants. Inducing ER stress in wild-type animals to levels equivalent to *asna-1(ok938)* mutants (**Fig. S2E**) and did not affect GFP::SEC-61β targeting (**Fig. 2C**). Starvation, which reduces insulin signaling and increases autophagy levels in *C. elegans* (Henderson and Johnson, 2001) and disrupts insulin secretion in *unc-31*/CAPS (Speese et al., 2007) mutants, did not affect GFP::SEC-61β targeting to the ER (**Fig. 2C**). Conversely, forcing high levels of DAF-28/insulin secretion in the *tom-1/tomosyn* mutant (Gracheva et al., 2007) failed to suppress the TAP targeting defect of *asna-1(ok938)* animals (**Fig. 2C**). Therefore, changes in ER morphology, insulin secretion activity, autophagy, or ER stress levels do not contribute to TAP targeting defects in *asna-1* mutants. These findings support the hypothesis that the insulin secretion function of ASNA-1 and high ER stress levels do not contribute to defective TAP targeting in *asna-1* mutants, thereby providing the first evidence of independent ASNA-1 functions in *C. elegans*.

### A receptor for TAP insertion interacts with ASNA-1 and has a role in cisplatin detoxification

ASNA-1-mediated insertion of TAPs requires a receptor at the ER membrane. WRB is a component of the heterodimeric receptor for TAP targeting (Vilardi et al., 2011). Its *C. elegans* homolog WRB-1 is the likely receptor for ASNA-1. Using co-IP/MS/MS analysis, we detected WRB-1 as an interaction partner of ASNA-1::GFP (**Table 1**). Western blot analysis showed a significant decrease in ASNA-1 protein levels in *wrb-1(tm5938)* mutants (**Fig. 2D**) indicative of their intracellular association. WRB-1::GFP was detected in the ER (**Fig. 2E**) and was required for GFP::SEC-61β targeting to the ER membrane (**Fig. 2A,B**) to roughly the same extent as ASNA-1. Furthermore, *wrb-1(tm5938)* produced a phenotype similar to *asna-1* mutants for proteinaceous aggregates (**Fig. S2A**), swollen ER (**Fig. S2B**), and autophagosomes (**Fig. S2C**). To determine whether WRB-1 is required for other ASNA-1 functions such as cisplatin responses, we tested the survival of *wrb-1(tm5938)* mutants after exposure to cisplatin; *wrb-1* mutants displayed cisplatin hypersensitivity (**Fig. 3D**). Thus, mutations in two different genes that similarly affect TAP targeting also demonstrated increased cisplatin sensitivity, suggesting that TAP targeting is associated with cisplatin sensitivity.

**Figure 3.**
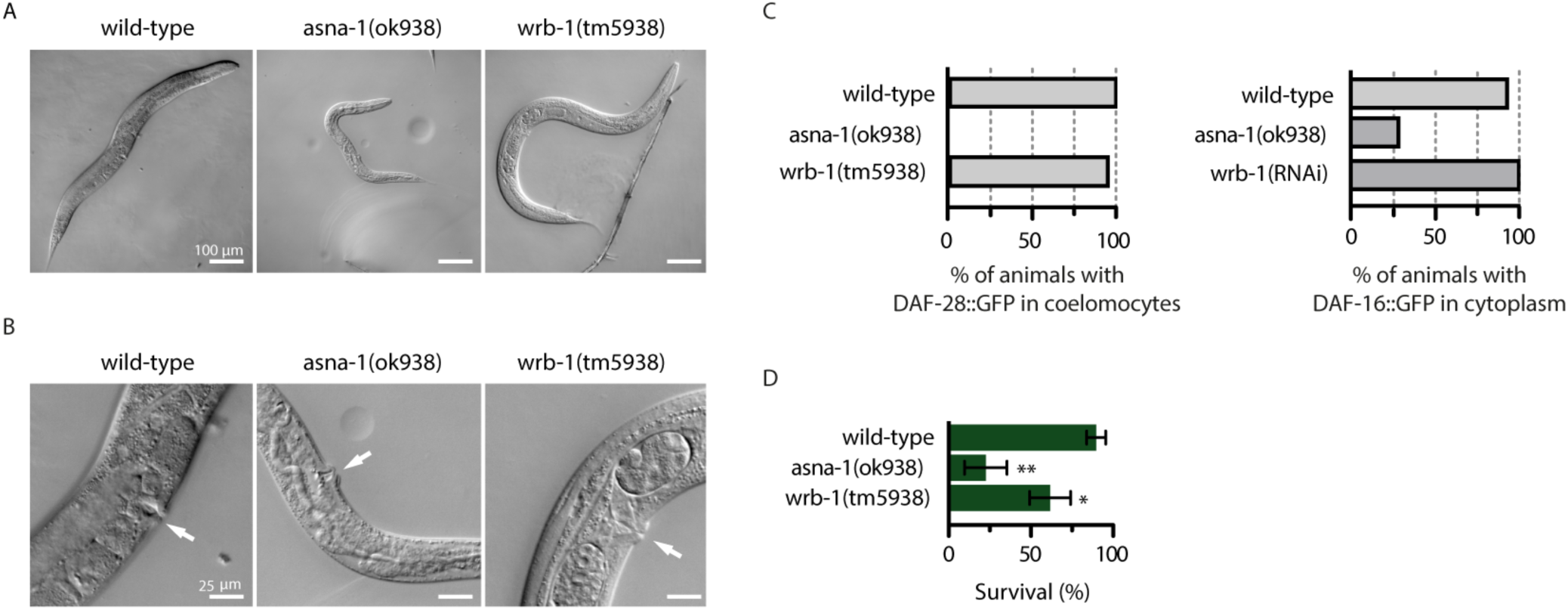
WRB-1 does not have a role in promoting insulin secretion. (A) Representative pictures of whole animals to reveal differences in body size between wild type, *asna-1(ok938)*, and *wrb-1(tm5938)* strains. (B) Magnification of the germline from animals shown in (A). White arrows indicate the position of the vulva. (C) Bar graph plot representing percentage of animals with secreted DAF-28:GFP in coelomocytes (n≥20) or the percentage of animals expressing DAF-16::GFP in the cytoplasm (n≥15). Error bars represent ± SD. (D) Bar graph plot of the mean survival rate of one-day-old young adult animals exposed to cisplatin for 24 h. Statistical significance was determined by the independent two-sample t-test (n ≥ 50, *p < 0.05, **p < 0.01).

### Cisplatin sensitivity is associated with TAP delocalization but not insulin secretion

We have previously shown that animals lacking *asna-1* activity reversibly arrest at the L1 stage and have defective insulin/IGF signaling (IIS). Furthermore, adult *asna-1(ok938)* mutants are pale, small, and sterile due to severe germline defects (Kao et al., 2007). While both ASNA-1 and WRB-1 mutants cause GFP::SEC-61β localization defects (**Fig. 2A,B**), *wrb-1*(*tm5938*) mutant adults are larger and have better developed germlines (**Fig. 3A,B**), indicating that TAP-targeting defects can be separated from the growth and body size phenotypes associated with insulin signaling defects. To directly determine whether WRB-1 loss leads to insulin secretion defects, we used two transgene reporters reporting IIS activity: DAF-16/FOXO::GFP (Henderson and Johnson, 2001) and DAF-28/insulin::GFP (Kao et al., 2007). In contrast to *asna-1(ok938)* mutants, *wrb-1(tm5938)* mutants had no DAF-28::GFP secretion defect (**Fig. 3C**), even though ASNA-1 protein levels were reduced (**Fig. 2D**). Furthermore, DAF-16/FOXO::GFP was always cytoplasmic in *wrb-1(RNAi)* animals, indicating high IIS levels (**Fig. 3D**). By contrast, DAF-16/FOXO:GFP localizes to the nuclei upon *asna-1* knockdown due to insulin signaling defects(Kao et al., 2007). *asna-1* (Kao et al., 2007) and insulin receptor mutants (Gems et al., 1998) exhibit reversible first larval stage arrest (**Fig. S3**). This characteristic insulin pathway growth defect was not observed when both maternal and zygotic *wrb-1*(*m-z-)* gene activity (**Fig. S3A, S3B**) were depleted. Taken together, these results show that, compared to mutants of its physical interaction partner ASNA-1, *wrb-1* mutants show no insulin secretion or signaling defects. Therefore, cisplatin sensitivity and TAP targeting defects are not associated with insulin secretion defects.

### Altered redox balance that favors ASNA-1^OX^ causes cisplatin sensitivity

To obtain direct evidence for the separable functions of ASNA-1, we tested a set of *asna-1* point mutant strains for cisplatin sensitivity (Thompson et al., 2013). *asna-1(gk592672)* substitutes a highly conserved alanine at position 63 to a valine, hereafter called *asna-1(A63V)*. This mutation did not affect the ASNA-1 protein stability (**Fig. 4A**). Transgene-expressed ASNA-1^A63V^::GFP protein was detected in the ER (**Fig. 4B**). The 13-times outcrossed *asna-1(A63V)* mutant was as sensitive to cisplatin exposure as the deletion mutant, and this phenotype was rescued by wild type transgene-expressed ASNA-1::GFP (**Fig. 4C**). This transgene also rescued the ASNA-1 protein null mutant for the cisplatin sensitivity phenotype (**Fig. 4C**). However, while as cisplatin sensitive as the deletion mutant, *asna-1(A63V)* animals had neither the insulin signaling/secretion defect (**Fig. 4F,G**) nor the enhanced ER stress phenotypes of the deletion mutant (**Figure S4A-C**). The animals were fertile with normal brood sizes (**Fig. S4E**), had properly developed germlines (**Fig. 4D,E**), and displayed a wild type lifespan (**Fig. S4D**). *asna-1(A63V)* animals had no TAP targeting defects (**Fig. 4H,I**), but membrane fractionation revealed reduced membrane association of the ASNA-1^A63V^::GFP protein compared to those with ASNA-1^+^:GFP (**Fig. 4J,K**). Further analysis revealed a strong defect in GFP::SEC-61β localization to the ER membrane in wild type animals as well as in *asna-1(A63V)* mutants when challenged with cisplatin (**Figure 4H,I**).

**Figure 4.**
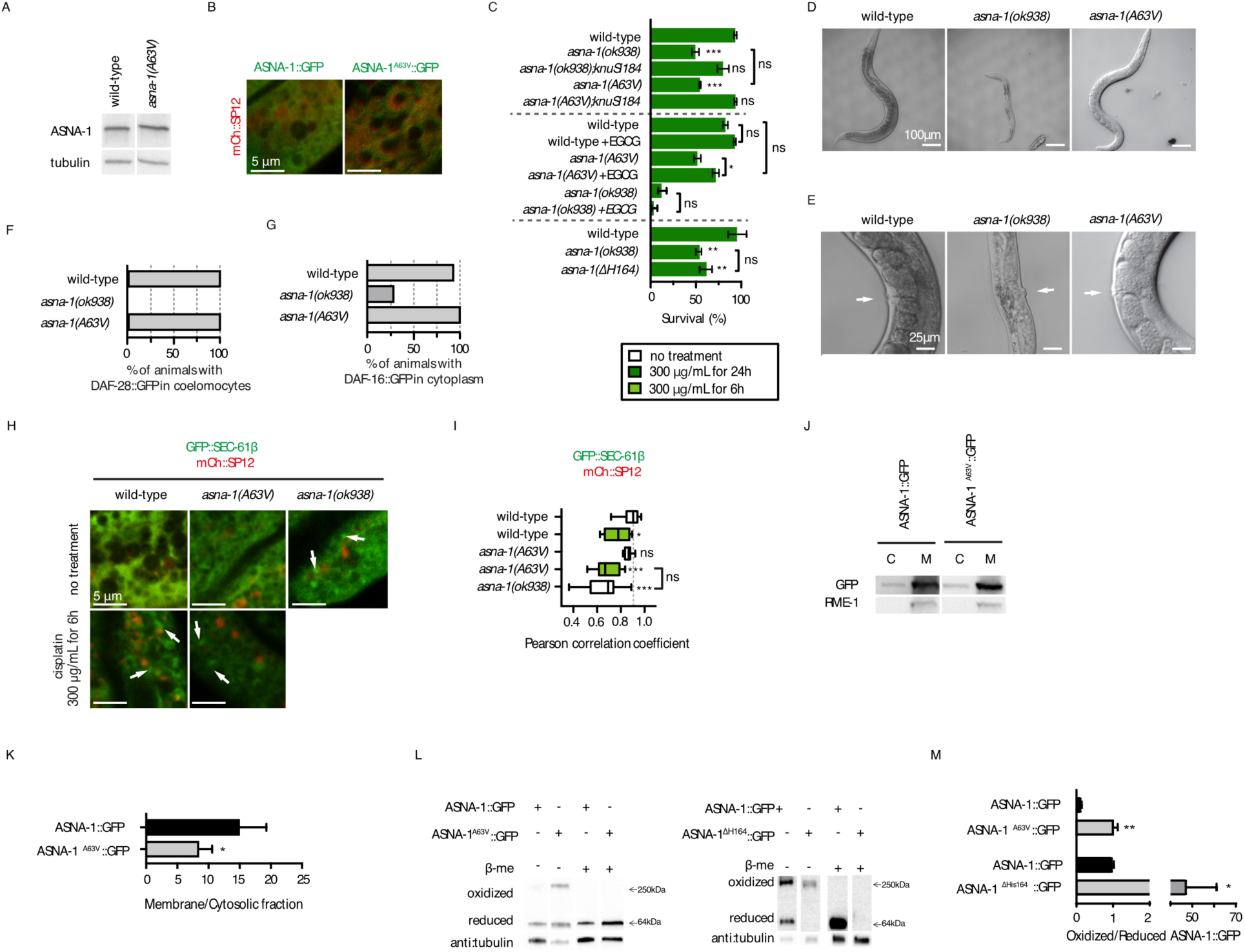
ASNA-1(A63V) increases cisplatin sensitivity via increased levels of oxidized ASNA-1. (A) Western blot analysis of ASNA-1 levels in wild-type and *asna-1(A63V)* animals. The blot was probed with anti-ASNA-1 antibody and tubulin was used as a loading control. (B) Confocal imaging merge of animals co-expressing ASNA-1::GFP or ASNA-1^A63V^::GFP with mCherry::SP12. (C) Bar graph plot representing mean survival ± SD of 48 h young adult animals after 24 h cisplatin exposure. EGCG pre-treatment: 400 μM for 48 h. Statistical significance was determined by independent two-sample t-test (n≥50). Error bars represent ± SD. (D) Representative pictures of animals/adult (hermaphrodites) worms revealing differences in body size between wild type, *asna-1(ok938)*, and *asna-1(A63V)* strains. (E) Magnification of the germline of the animals from (A). White arrows indicate the vulva position. (F) Bar graph plot representing percentage of animals with secreted DAF-28:GFP in coelomocytes (n ≥ 20). Error bars represent ± SD. (G) Bars graph plot representing percentage of animals expressing DAF-16::GFP in cytoplasm (n ≥ 15). Error bars represent ± SD. (H) Confocal imaging merge of wild-type, *asna-1(A63V)*, and *asna-1(ok938)* animals co-expressing GFP::SEC-61β and mCherry::SP12 with or without cisplatin treatment (300 μg/mL for 6 h). (I) Pearson correlation analysis of GFP::SEC-61β and mCherry::SP12 colocalization in different strains. Box plot represents the average Pearson correlation coefficient (R) of indicated strains. Statistical significance was determined by one-way ANOVA followed by Bonferroni post-hoc correction (n≥10; *p<0.05, **p<0.01, ***p<0.001). (J) Subcellular localization of ASNA-1::GFP and ASNA-1^A63V^::GFP. C - cytosolic fraction; M - membrane fraction. Blot was probed with anti-GFP antibody. Anti-RME-1 antibody was used as a marker for membrane proteins. (K) Band intensity quantification of membrane/cytosolic fraction of ASNA-1::GFP and ASNA-1^A63V^::GFP. Statistical significance was determined by the independent two-sample t-test. Error bars represent ± SD. (L) Reducing and non-reducing SDS-PAGE visualizes levels of oxidized and reduced ASNA-1^A63V^::GFP and an ASNA-1^ΔHis164^::GFP-expressing transgenic line. Control refers to worms expressing the wild-type ASNA-1::GFP transgene. Blots were probed with anti-GFP antibody and tubulin was used as a loading control. In each gel the figure is composed of lanes from a single gel. (M) Band intensity quantification of oxidized/reduced ASNA-1::GFP in different transgenic lines from the experiments in (L). Statistical significance was determined by the independent two-sample t-test. Experiments were performed in triplicate. Error bars represent ± SD.

The antioxidant epigallocatechin gallate (EGCG) reduces cisplatin effectiveness (Pan et al., 2015) and has been used as an antioxidant in several *C. elegans* studies (Xiong et al., 2018; Zhang et al., 2009). Notably, pre-treatment with EGCG reduced *asna-1(A63V)* cisplatin sensitivity but had no protective effect in the *asna-1* deletion mutant (**Fig. 4C**). Modulation of cisplatin sensitivity by EGCG suggested that the effect of the A63V change might alter the balance between oxidized and reduced ASNA-1 and that ASNA-1 was a direct target for the antioxidant effect of EGCG. Comparison of oxidized and reduced ASNA-1^A63V^::GFP to ASNA-1::GFP revealed that the point mutation shifted the balance from the reduced to the oxidized form (**Fig. 4L,M**), indicating that a shift to more oxidized ASNA-1 caused increased cisplatin sensitivity. To examine this further, we tested the redox balance in worms expressing ASNA-1^ΔHis164^::GFP, which is inactive for rescue of the cisplatin response phenotype (Hemmingsson et al., 2010). Again, substantially more oxidized ASNA-1 was detected (**Fig. 4L,M**). To examine this further, we tested an *asna-1(ΔHis164)* mutant strain generated by Crispr/CAS9 technology. This strain was as sensitive to cisplatin as the *asna-1(ok938)* deletion mutant (**Fig. 4C**). Thus, two ASNA-1 mutants that displayed increased oxidation were as sensitive to cisplatin as the deletion mutants. Moreover, *sod-2(gk257)* and *mev-1(kn1)* mutants, which harbor high levels of ASNA-1^OX^ without changes in overall ASNA-1 levels (**Fig. 1C,D**), were also cisplatin sensitive (**Fig. 6C**). Taken together, a redox balance that favors oxidized ASNA-1 results in the same cisplatin sensitivity phenotype as complete depletion of ASNA-1, leading to the conclusion that reduced ASNA-1 protects against cisplatin toxicity. Notably, cisplatin sensitivity becomes a redox-sensitive event in the *asna-1^A63V^* genetic background, indicating a strategy to modulate its killing effect. Strikingly, the shift in ASNA-1^A63V^ to the oxidized state occurred even in the absence of an increase in the expression of markers of ER stress (*hsp-4*) (**Fig. S4A-C**), mitochondrial stress (*hsp-6* and *hsp-60*) (**Fig. S4F**), or oxidative stress (*gst-4*, *gst-30* and *gst-38*) (**Fig. S4G**) in contrast to high ROS levels in *sod-2(gk257)* and *mev-1(kn1)* mutants. We conclude that the A63V and ΔHis164 mutants in ASNA-1, which have inherently high ASNA-1^OX^ levels, are sensitive to cisplatin but have normal insulin secretion function. This analysis proves that the redox balance of ASNA-1 is necessary for maintaining its essential functions and that reduced ASNA-1 protects against cisplatin toxicity, most likely through its TAP targeting activity.

### Cisplatin inactivates the protective function of ASNA-1 by converting it to the oxidized state

Having shown that reduced ASNA-1 is protective, we sought to examine whether part of cisplatin’s killing mechanism was by directly targeting and inactivating ASNA-1. Several groups have shown in mammalian cancer cell models that a central feature of cisplatin treatment is oxidative stress induction and ROS generation (Brozovic et al., 2010; Choi et al., 2015; Itoh et al., 2011; Lu et al., 2016). Since ASNA-1 is required for cisplatin resistance and ASNA-1 function is modulated by oxidation, we next asked if cisplatin exposure induced oxidative stress in *C. elegans.* A short cisplatin exposure regimen was chosen at 300 µg/mL or 600 µg/mL, since these concentrations do not decrease the viability of wild-type animals, thereby allowing meaningful cell biology analysis. These experimental conditions were indeed sufficient to induce significant elevations in ROS (**Fig. 5D**), increased protein carbonylation (**Fig. 5C**), and robustly activate the oxidative stress response genes *gcs-1* and *gst-4* (**Fig. 5A,B**). We next tested whether cisplatin exposure influences the oxidation state of ASNA-1. ASNA-1::GFP-expressing worms were exposed to cisplatin at a concentration that significantly reduced the survival of *asna-1(ok938)* mutants without affecting wild-type animals (**Fig. 4C**). Under these conditions, the balance of ASNA-1::GFP shifted towards the oxidized form (**Fig. 5E,F**). Thus, cisplatin exposure directly increases *asna*-*1* oxidation, allowing us to conclude that ASNA-1 is a molecular target of cisplatin.

**Figure 5.**
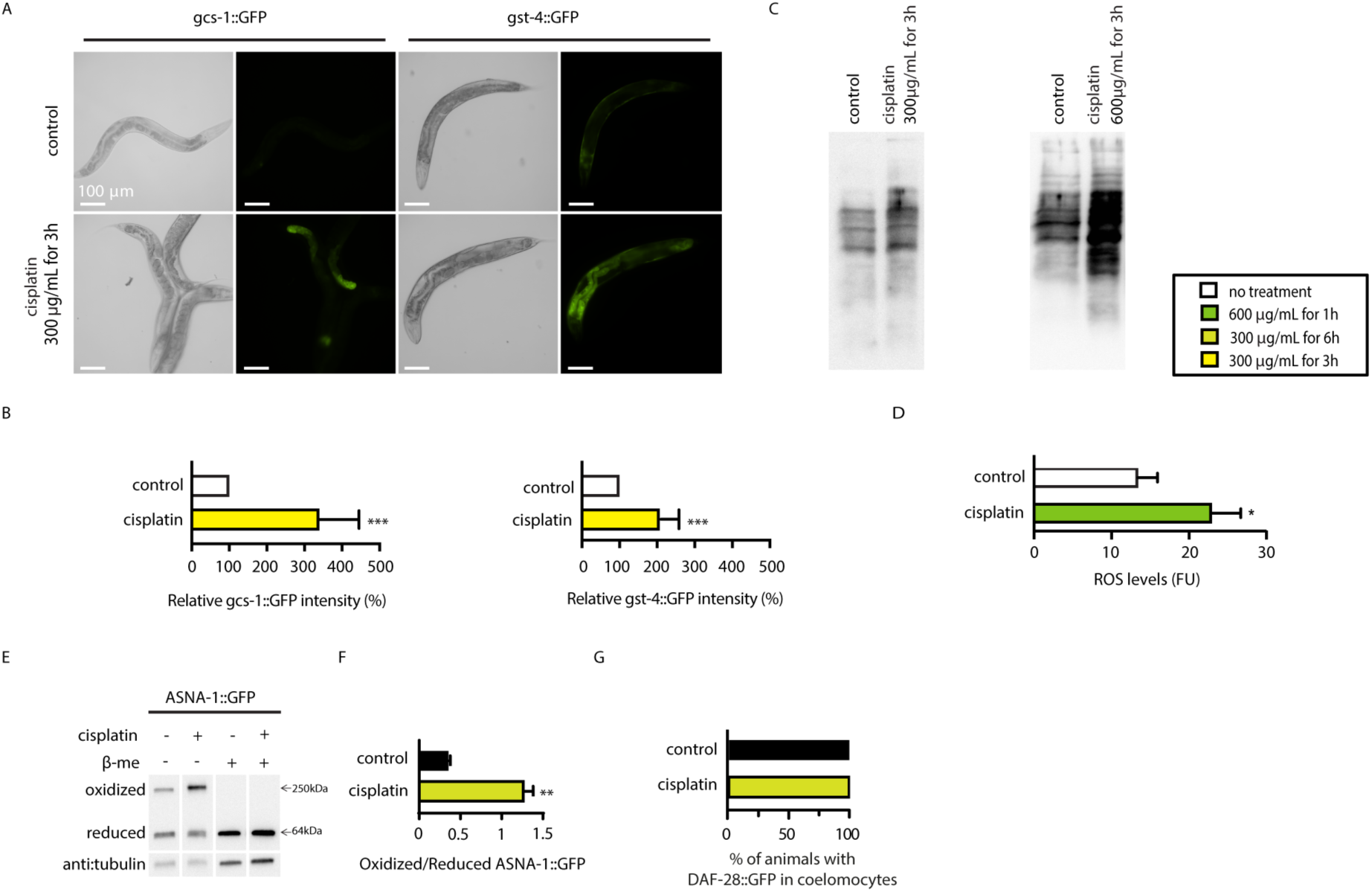
Cisplatin directly increases ROS levels and oxidation of ASNA-1 without affecting insulin secretion. (A) Increase of GFP intensity upon cisplatin exposure (300 µg/mL for 3 h) in *gcs-1::GFP* and *gst-4::GFP* transgenic strains. (B) Quantification of relative GFP intensity for *gcs-1::GFP* (n ≥ 15) and *gst-4::GFP* (n ≥15) transgenic strains. Statistical significance was determined by the independent two-sample t-test (*p < 0.05, **p < 0.01, ***p < 0.001). Bars represent mean ± SD. (C) Influence of cisplatin exposure on protein oxidation evaluated using the OxyBlot kit. Proteins recovered from lysed worms exposed to cisplatin for 3 h at a concentration of 300 μg/mL or 600 μg/mL were labeled with DNP solution (Oxyblot) to visualize the presence of protein carbonyls by immunoblotting. (D) ROS production levels as estimated by the H_2_DCFDA assay. Wild-type strain was exposed to 600 μg/mL cisplatin for 1 h. Statistical significance was determined by the independent two-sample t-test (n ≥ 1000). Experiments were performed in triplicate (*p < 0.05). Bars represent mean ± SD. (E) Reducing and non-reducing SDS-PAGE visualizes oxidized and reduced ASNA-1::GFP exposed to cisplatin (300 µg/ml for 6 h). Blot was probed with anti-GFP antibody and tubulin was used as a loading control. The figure is composed of lanes from a single gel. (F) Band intensity quantification of oxidized/reduced ASNA-1::GFP exposed to cisplatin (300 µg/ml for 6 h). Statistical significance was determined by the independent two-sample t-test. Error bars represent ± SD. (G) Bars represent percentage of animals with secreted DAF-28:GFP in coelomocytes (n≥20) exposed to cisplatin (300µg/ml for 6 h). Error bars represent ± SD.

**Figure 6.**
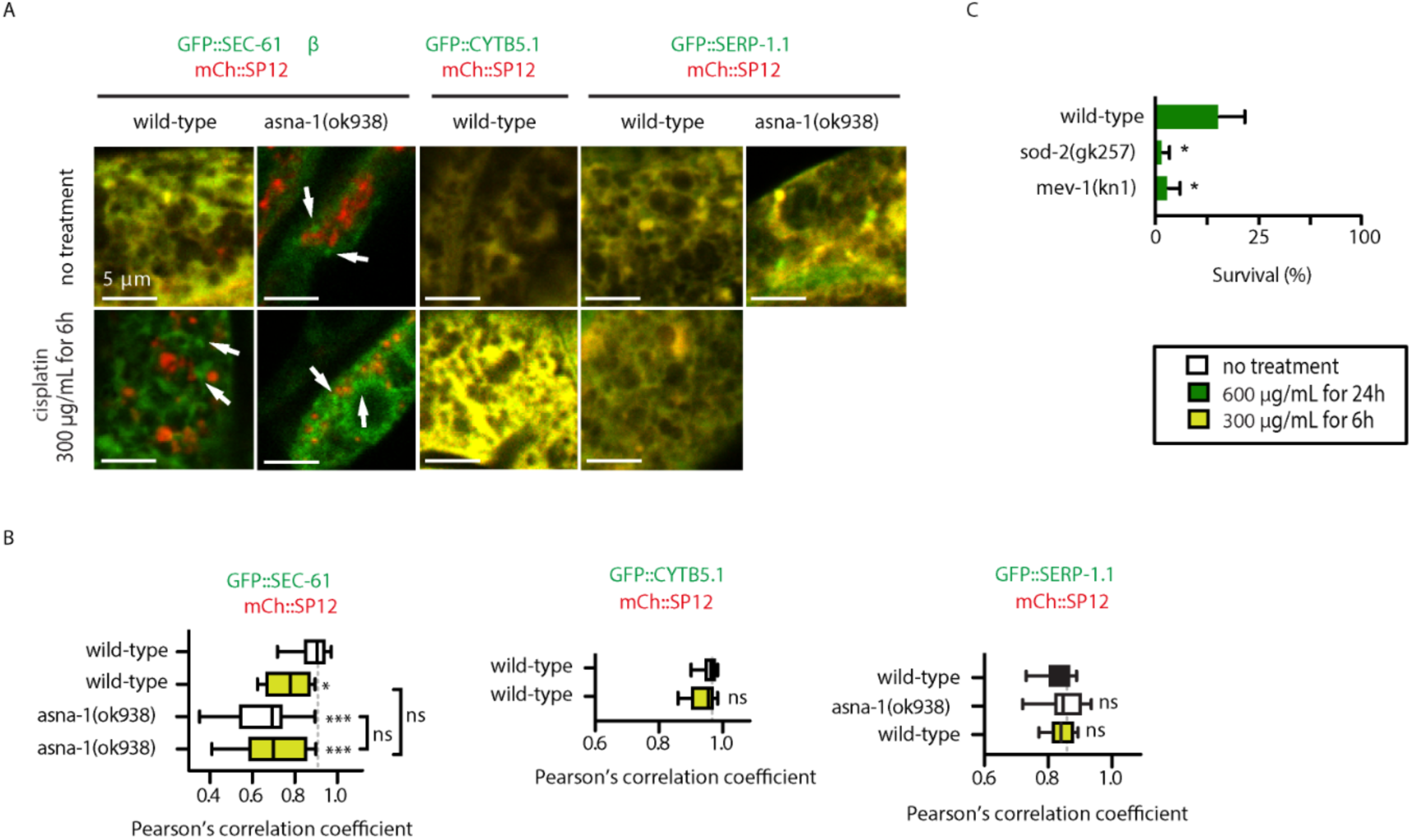
Cisplatin selectively delocalizes ASNA-1-dependent TA protein. (A) Confocal imaging merge of wild-type or *asna-1(ok938)* animals co-expressing GFP::SEC-61β, GFP::CYTB5.1, or GFP::SERP-1.1 with mCherry::SP12, with or without cisplatin treatment (300 μg/mL for 6 h). (B) Pearson correlation analysis of GFP::SEC-61β, GFP::CYTB5.1, or GFP::SERP-1.1 and mCherry::SP12 co-localization. Box plot represent the average Pearson correlation coefficient (R) of the indicated strains without or with cisplatin treatment (300 μg/mL for 6 h). Statistical significance was determined by one-way ANOVA followed by Bonferroni post-hoc correction (n≥10). (*p<0.05, **p<0.01, ***p<0.001). (C) Bars represent mean survival ± SD of one-day-old young adult animals exposed to cisplatin. Statistical significance was determined by the independent two-sample t-test (n≥50). Experiments were performed in triplicate.

### Cisplatin selectively delocalizes an ASNA-1-dependent TAP from the ER

Having established that ASNA-1^RED^ protects against cisplatin toxicity and mindful of the role of the reduced form in TAP targeting, we next determined whether cisplatin treatment and TAP targeting were directly related. To this end, we tested whether the increased oxidized ASNA-1 observed in cisplatin-treated worms resulted in a biologically meaningful outcome. Exposure of *asna-1(+)* animals expressing GFP::SEC61β to cisplatin significantly delocalized this TAP away from the ER membrane at levels similar to those seen in untreated *asna-1(ok938)* animals (**Fig. 6A,B**). Remarkably, cisplatin treatment had no effect on the ER localization of the two ASNA-1-independent TAPs CytB5.1 (*cytb-5.1)* and SERP-1.1 (**Fig. 6A,B**). Quantitative confocal microscopy revealed that conditions that substantially delocalized GFP::SEC-61β had no effect on the ER localization of GFP::CYTB-5.1 and GFP::SERP-1.1. Therefore, delocalization of GFP::SEC-61β in cisplatin-treated worms was not due to more a more generalized effect on membrane integrity. We next tested the effect of cisplatin exposure on DAF-28/insulin::GFP secretion and observed no secretion defect (**Fig. 5G**); the membrane events associated with insulin maturation, packaging into dense core vesicles, and release were unaffected. Therefore, the effects on GFP::SEC-61β localization were not caused by general cisplatin effects on cellular function and animal health. Since cisplatin oxidizes ASNA-1 and shifts the protein to its non-TAP associated role, we next asked whether GFP::SEC-61β delocalization in cisplatin-treated animals was solely via its modulation of ASNA-1 activity or due to another role in this process. ER targeting of GFP::SEC61β was compared in *asna-1(ok938)* mutants with and without cisplatin treatment; GFP::SEC61β was delocalized to the same extent in both conditions (**Fig. 6A,B**). Since there was no additive effect, cisplatin exposure requires ASNA-1 activity to exert its TAP delocalization effect.

Our studies support the hypothesis that cisplatin directly targets ASNA-1, leading to its rapid oxidation without affecting insulin secretion. We conclude that a shift in redox balance that lowers ASNA-1^RED^ levels blocks insertion of its client TAP into the ER membrane. Furthermore, cisplatin treatment of wild-type worms phenocopied the *asna-1* deletion mutant phenotype for delocalization of GFP::SEC61β but not for the insulin secretion function. The ER is an organelle whose function is affected by cisplatin exposure via its effect on ASNA-1 oxidation.

## DISCUSSION

Cisplatin-generated ROS is a central event in post-mitotic cells. The precise targets of these oxygen species are poorly understood, and their identification remains critical to understanding cisplatin cytotoxicity. Here we identified ASNA-1 as an oxidative stress-induced molecular target of cisplatin and provide a molecular basis for the cisplatin hypersensitivity caused by ASNA-1 knockdown.

This study was underpinned by the observations that cisplatin induces oxidative stress (Itoh et al., 2011), ASNA-1 knockdown increases sensitivity to cisplatin (Hemmingsson et al., 2010), that the yeast homolog, GET3, exists in both oxidized and reduced states, and that GET3 must be reduced to target some TAPs to the ER membrane (Voth et al., 2014). Using live cell imaging, we have shown that worm ASNA-1 also has TAP targeting activity and that WRB-1 is its likely receptor for this function. *wrb-1* mutants were also cisplatin sensitive, thereby linking TAP targeting to cisplatin sensitivity and uncovering a new cisplatin sensitivity factor. ASNA-1 existed in both oxidized and reduced states in a manner requiring redox-reactive cysteines. We provide several lines of evidence showing that elevated ASNA-1^OX^ levels increases cisplatin sensitivity. Strikingly, a redox-imbalanced mutant that favored accumulation of ASNA-1^OX^ was as sensitive as a mutant lacking ASNA-1. Cisplatin exposure rapidly increased ROS and ASNA-1^OX^ levels. Moreover, an ASNA-1-dependent TAP was specifically delocalized as a result of cisplatin treatment.

### ASNA-1 is required for a TAP-targeting protein

We characterized the TAP-targeting activity of *C. elegans* ASNA-1 and found that it possesses robust TAP targeting activity shared with its binding partner, the ER-localized protein WRB-1. *C. elegans asna-1* has all the structural features required for TAP insertion (Chio et al., 2019)(Mateja and Keenan, 2018; Mateja et al., 2009). However, since not all ASNA-1 homologs have TAP-targeting activity (Farkas et al., 2019), it was important to examine whether *C. elegans* ASNA-1 has this function. In addition, parallel pathways like EMC, HSP70/HSC40, and SND participate in TAP targeting (Aviram et al., 2016; Casson et al., 2017; Cho and Shan, 2018; Guna et al., 2018). It was possible that compensated TAP targeting by these pathways would make it difficult to detect a targeting defect in *asna-1* single mutants. Consequently, it was important to establish the *in vivo* contribution of ASNA-1 to TAP targeting in intact animals. We assayed this activity in large intestinal cells, because ASNA-1 expressed by these cells was amenable to quantitative confocal microscopy. We showed that the other pathways do not bypass the need for ASNA-1 activity, since SEC-61β, which binds to ASNA-1 (**Table 1**), was significantly delocalized in *asna-1* mutants (**Fig. 2A,B**). Importantly, using this assay, we also found two ASNA-1-independent ER-localized TAPs (**Fig. 6A**), in agreement with findings in other systems (Colombo et al., 2009; Favaloro et al., 2008; Stefanovic and Hegde, 2007). The contribution of other pathways to the targeting of ASNA-1-dependent TAPs in worms is unclear. It is possible that if TAP targeting is studied in *C. elegans* tissues other than the intestines, the EMC or the HSP70/HSC40 pathways may play a greater role in the process.

### WRB-1 participates in TAP targeting and is the likely ER receptor for ASNA-1

*C. elegans* WRB-1 was identified as the closest homolog of mammalian WRB, containing two predicted trans-membrane domains and the important coiled-coil domain. Evidence that WRB-1 is the receptor for the ASNA-1/TAP complex comes from the fact that it was ER-localized (**Fig. 2E**) and was an ASNA-1-binding protein (**Table 1**), as with its homologs in other species (Carvalho et al., 2019; Vilardi et al., 2011). Moreover, ASNA-1 protein levels were lower in *wrb-1* mutants (**Fig. 2D**), a property also seen upon mammalian WRB knockdown (Colombo et al., 2016; Rivera-Monroy et al., 2016). Taken together, these three properties of WRB-1 satisfy the requirement to regard this protein as the ER-based receptor. Crucially, TAP localization was defective in two *wrb-1* mutants to a similar extent seen in *asna-1* mutants (**Fig. 2B**).

### Cisplatin cytotoxicity is linked to defects in TAP targeting but not the insulin secretion function of ASNA-1

The observation that TAP targeting was defective in *wrb-1* mutants allowed us to address two important questions about TAP protein function. First, we showed that *wrb-1* mutants were also hypersensitive to cisplatin (**Fig. 3D**), thus identifying mutants in two separate genes with TAP targeting and cisplatin hypersensitivity function. Second, we found that *wrb-1* mutants were not defective in insulin/IIS signaling and secretion function (**Fig. 3**), as observed in *asna-1* mutants (Kao et al., 2007). We therefore concluded that insulin/IIS signaling pathway defects play no role in the *asna-1(−)* cisplatin hypersensitivity phenotype. Moreover, this also meant that ASNA-1-dependent TAP targeting has little role in ASNA-1-induced DAF-28/insulin secretion. This was consistent with our previous finding that *daf-2/insulin* receptor mutants without insulin signaling capacity are completely resistant to cisplatin (Hemmingsson et al., 2010). Further evidence that the cisplatin and insulin functions of ASNA-1 are separable emerged from the analysis of the *asna-1(A63V)* mutant, which had the striking phenotype of completely separating the cisplatin and insulin secretion functions (**Fig. 4**): “null” for the cisplatin phenotype and “wild-type” for the growth and insulin secretion function of ASNA-1.

Since ASNA-1 protein levels were similar in wild-type and the *asna-1(A63V)* mutant (**Fig. 4A**), this functional separability (cisplatin versus insulin secretion) was not due to a threshold effect, in which a differential phenotypic effect would be seen if one function required less total ASNA-1 protein.

### *C. elegans* ASNA-1 is a redox-sensitive protein

*C. elegans* ASNA-1 exists in both reduced (ASNA-1^RED^) and oxidized (ASNA-1^OX^) forms (**Fig. 1, Fig. 7A**). This is consistent with Voth *et al*. (2014), who showed that yeast GET3 changes from a reduced, ATPase-dependent TAP targeting state to an oxidized ATPase-independent general chaperone/holdase state. The ASNA-1^RED^/ASNA-1^OX^ balance was biologically relevant, since it was sensitive to changes in animal physiology. Worm mutants with high ROS levels or cultivation conditions that increase ROS shifted the balance towards more oxidized ASNA-1, while exposure to antioxidants shifted the balance in the opposite direction (**Fig. 1**). The ASNA-1^RED^/ASNA-1^OX^ balance had biological relevance, since cisplatin sensitivity was increased in the *asna-1(A63V)* mutant that produced a preferentially oxidized protein variant (**Fig. 4L, Fig. 7B**). Notably, the cisplatin hypersensitivity of *asna-1(A63V)* animals was reduced by pre-treatment with the antioxidant EGCG, indicating that the redox state of ASNA-1(A63V) is relevant for cisplatin cytotoxicity (**Fig. 4C**). Reversal of the cisplatin sensitivity phenotype with antioxidants strongly supports this conclusion. The shift in the ASNA-1^RED^/ASNA-1^OX^ balance towards higher ASNA-1^OX^ levels is not a peculiarity of the alanine 63 mutation; indeed, ASNA-1^ΔHis164^ displayed even higher ASNA-1^OX^ levels (**Fig. 4L**). Transgenic expression of ASNA-1^ΔHis164^ does not rescue cisplatin hypersensitivity of the null mutant but maintains largely intact insulin/IGF signaling (Hemmingsson et al., 2010). We speculate that ASNA-1 point mutants that are TAP-targeting defective exist at higher levels in the oxidized state.

**Figure 7.**
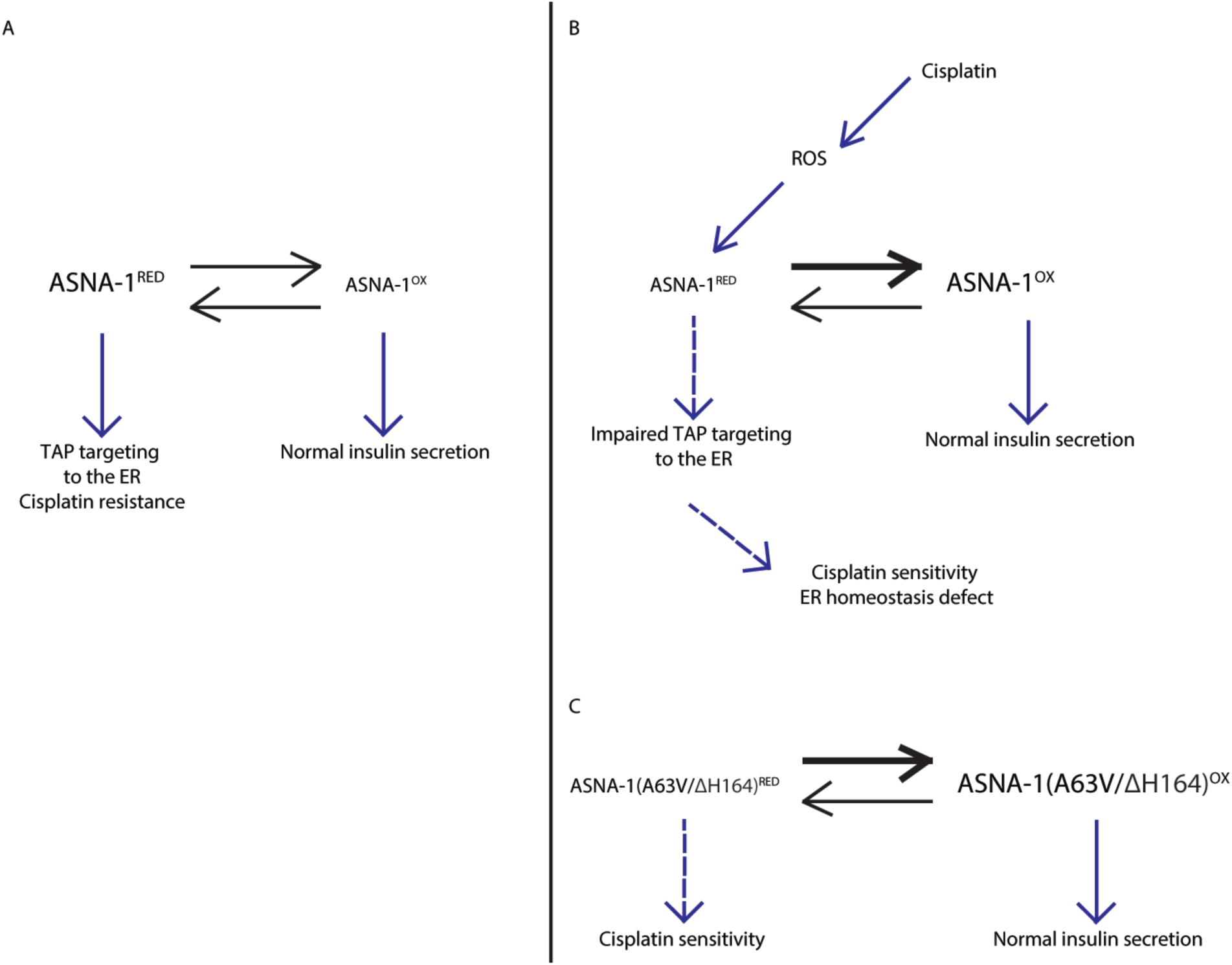
ASNA-1-mediated roles in cellular function and cisplatin sensitivity are based on the balance of its redox states. (A) Under normal physiological conditions, ASNA-1 is present in both forms: reduced (ASNA-1^RED^) and oxidized (ASNA-1^OX^). ASNA-1^RED^ participates in the insertion of TAP into the ER membrane, while ASNA-1^OX^ assures proper insulin secretion. (B) When the cells are exposed to cisplatin, elevated ROS levels perturb the ASNA-1 redox balance. ASNA-1 shifts towards the ASNA-1^OX^ state at the expense of ASNA-1RED, which impairs TAP targeting to the ER without affecting insulin secretion. Because less ASNA-1^RED^ is available, ER homeostasis is affected and cells are more sensitive to cisplatin. (C) Both ASNA-1-mutant proteins (A63V and ΔH164) are found predominantly in the ASNA-1^OX^ state, which results in greater cisplatin sensitivity but unaffected insulin secretion.

ASNA-1 is member of a group of *C. elegans* proteins, along with PRDX-2, IRE-1, and GLB-12, whose activity changes with oxidation (Henau et al., 2015; Hourihan et al., 2016; Oláhová et al., 2008). *In vivo* redox changes are dynamic in *C. elegans* with respect to both tissue type and age (Back et al., 2012; David et al., 2010; Walther et al., 2015). Oxidized proteins are likely to be widespread in the proteome, since quantitative redox proteomics has identified many proteins with redox-sensitive cysteines in H_2_O_2_-treated animals (Kumsta et al., 2011) and in day 2 adults (Knoefler et al., 2012).

### Increased levels of oxidized ASNA-1 sensitize cells to cisplatin killing

Only 5-10% of cisplatin is found in the nuclei of exposed cells, the remainder being cytoplasmic and membrane associated (Brozovic et al., 2010). Non-nuclear cisplatin has been shown to modify kinase signaling, Ca^2^+ pump activity, and membrane dynamics (Brozovic et al., 2010; Dasari and Tchounwou, 2014). In light of our findings that cisplatin modifies ASNA-1 function (**Fig. 7B**), it will be instructive to study which - if any - of these processes are caused by changes in ASNA-1 oxidation levels and ASNA-1-dependent TAP targeting. Our study shows that cisplatin has an effect on the ER in worm intestinal cells, which was not due to a general effect on cellular membranes at the concentrations of cisplatin used, since only ASNA-1-dependent TAPs were delocalized and ASNA-1-independent TAPs were targeted normally (**Fig. 6**). Further, this treatment had no effect on DAF-28/insulin secretion, a process that also requires several membrane budding and fusion events at various cellular locations (**Fig. 5G**). The normal insulin secretion levels detected in cisplatin-treated animals suggest that Golgi and plasma membranes are largely competent.

Most cells in human solid tumors are in a post-mitotic state (Komlodi-Pasztor et al., 2012), similar to in adult worms. Given the range of effects exerted by cisplatin on cellular function, the killing effect cannot only be attributed to induced DNA damage but is also likely to be partially be due to effects on the ER. We have shown that increased levels of oxidized ASNA-1 beyond a certain threshold will sensitize cells to cisplatin. We propose that small molecule drugs that increase oxidized ASNA-1 at the expense of the reduced form will enhance cisplatin cytotoxicity. Conversely, the antioxidant EGCG reduces cisplatin effectiveness. This work also reveals for the first time that cisplatin impacts TAP targeting. The cellular biology of the TAP pathway has been studied in detail and the roles of several other proteins besides ASNA1/TRC40 delineated. It is likely that small molecule drugs that affect other components of the TAP targeting pathway would also increase cisplatin sensitivity; indeed, *wrb-1* mutants were as sensitive as *asna-1* mutants and also represent a new cisplatin sensitivity factor. This analysis of ASNA-1 demonstrates that cisplatin perturbs ER function, which might also explain other effects of cisplatin on signaling pathways, Ca^2+^ pump function, and membrane properties. We previously showed that while cisplatin does not induce ER stress, combinatorial use with an ER stress inducer sensitizes resistant worms to cisplatin (Natarajan et al., 2013). This is consistent with the idea that cisplatin treatment sensitizes the ER in a metastable state due to TAP targeting defects and if ER function is further compromised then the cells will become hypersensitive to cisplatin cytotoxicity. Further, increased endogenous levels of oxidized ASNA-1 increases cisplatin sensitivity. Drugs that increase ASNA-1 oxidation, target other components of the TAP biogenesis pathway, or induce mild effects on ER function will enhance cisplatin sensitivity, address the problem of cisplatin resistance, and permit lower combinatorial doses. Modeling drug action in non-mammalian genetic model systems is extremely useful for identifying potential protein targets to improve cancer therapy.

## Material and Methods

### Plasmids

pVB343ML(Larsen et al., 2007): 1.8 kb of the intestine-specific *vha-6* promoter was cloned as a NheI/SacI fragment into pPD49.26 (Fire lab vector kit). This was used as a backbone for all subsequent cloning with fluorescent markers. pVB637OB and pVB638OB:: GFP or mCherry was amplified without stop codons and inserted as KpnI/NheI fragments after the *vha-6* promoter in pVB343ML. GFP insertion produced pVB637OB and mCherry produced pVB638OB. pVB641OB: Genomic *spcs-1* (SP12) was amplified and inserted as a KpnI/KpnI fragment after and in frame with the mCherry in pVB638OB. pVB643GK: The entire cDNA for *wrb-1* (Y50D4A.2) was amplified as a HinDIII/HindIII fragment and sub-cloned into the TOPO TA vector. From there it was excised with HindIII and cloned into the L4440 to give pVB643GK. This was the template used to make *wrb-1* dsRNA with the MegaScript T7 kit (ThermoScientific). pVB644OB: The full-length cDNA of *wrb-1* was synthesized as a KpnI/KpnI fragment (GenScript Inc., NJ, USA) and cloned 3’ to, and in frame with the GFP in pVB637OB. The entire construct was sequenced and found to be error-free.

GFP::SEC-61 β: The open reading frame of Y38F2AR.9 *(sec-61β)* was amplified and inserted as a KpnI/KpnI fragment after and in frame with the GFP in pVB637OB to yield pVB639OB. GFP::CYTB-5.1 The open reading frame of *cytb-5.1* (Cytochrome B5) was amplified and inserted as a KpnI/KpnI fragment after and in frame with the GFP in pVB637OB to yield pVB640OB. GFP::SERP-1.1: The genomic region of the TAP *serp-1.1* lacking the initiator methionine was amplified with Q5 polymerase and inserted downstream and in frame with GFP in pVB637OB to generate the plasmid pDR28. GFP: SEC-61β lacking the transmembrane domain: The genomic clone for worm sec-61β lacking the TMD was synthesized by Genscript (Piscataway, USA), and the clone lacking the initiator methionine was inserted in frame downstream of GFP in pVB637OB using Gibson ligation technology to yield pGK208.

### *C. elegans* genetics and maintenance

The Bristol strain (*N2*) was used as wild-type control and served as parental line for all subsequent strain constructions. N2, *zIs356(daf-16::gfp)* and *svIs69(daf-28::gfp)* are described in WormBase (www.wormbase.org). *wrb-1(tm5938 & tm5532)* were obtained from the the Mitani lab and NBRP, Tokyo (www.shigen.nig.ac.jp/c.elegans/index.jsp). *wrb-1(tm5938)* was outcrossed six times to N2 and maintained in trans to the *nT1(qIs51)* balancer. *asna-1(ok938)/hT2(qIs48)* was maintained as previously described (Kao et al., 2007). The *asna-1(gk592672)* containing strain VC40357 was outcrossed 13 times using *unc-32(e189)* and *oxTi719* [*eft-3p::tdTomato::H2B::unc-54’UTR Cbr-unc-119(+)*] as balancers and kept *in trans* to the *hT2(qIs48)* balancer. *svIs135* [*Pvha-6::gfp::sec-61β(Y38F2AR.9)+Pvha-6::mCherry::SP12*] was generated by microinjection of pVB639OB together with pVB641OB at 50 ng/μl each into wild-type N2, to generate an extrachromosomal array followed by its genomic integration using gamma irradiation. Integration of the *svIs135* transgene was on the X chromosome. *svIs135* bearing worms were backcrossed six times to N2 to eliminate background mutations. *svIs135;asna-1(ok938)/hT2(qIs48)*, *svIs135;asna-1(A63V)/hT2(qIs48)* and *svIs135;wrb-1(tm5938)/nT1(qIs51)* strains were generated using the 6x outcrossed version of *svIs135. svIs143(Pnhx-2::mCherry::lgg-1)* was generated by genomic integration (as above) of *vkEx1093*(Gosai et al., 2010) and outcrossed three times to N2. The 3x outcrossed version of *svIs143* was used to generate *svIs143; asna-1(ok938)/hT2(qIs48)*. svEx917 (*Pvha-6::gfp::cytb-5.1; Pvha-6::mCh::SP12*) was generated by injecting pVB640OB and pVB641OB at 50 ng/μl each into wild-type N2. Injection RNAi of *wrb-1* dsRNA (made using pVB643GK as a template). RNAi was performed as previously described (Billing et al., 2012) ASNA-1::GFP(*svIs56*) has been previously described (Kao et al., 2007). ASNA-1^A63V^::GFP was generated by mutagenesis of pVB222GK (Kao et al., 2007) as a template to introduce the A63V change by the QuickChange II Site-Directed Mutagenesis Kit (Agilent Technologies). The ASNA-1^A63V^::GFP containing plasmid (pGK200) was used for generating the extrachromosomal arrays *rawEx8* and *rawEx9. rawEx8* bearing worms were irradiated for chromosomal integration to yield the *rawIs16* transgene. Animals bearing *rawIs16* were outcrossed 4 times to N2 before analysis. The integrated transgene *rawIs13* that expresses *asna-1^C285S;C288S^::GFP* was obtained by gamma ray irradiation of worms bearing the *svEx756* extrachromosomal transgene (Hemmingsson et al., 2010). The single copy *asna-1::gfp(knuSi184)* consisted of a 1.4 kb upstream promoter sequence driving genomic *asna-1* coding region to which the GFP coding region was fused just before the stop codon. *tbb-2* was used as the 3’UTR for this transgene. This was inserted on chromosome II at the *ttTi5605* locus using MosSCI technology in the *unc-119(ed3)* background (Frøkjær-Jensen et al., 2008) by Knudra Transgenics. *sod-2(gk257*), *mev-1(kn-1*), *hsp-4::GFP (zcIs4*), *unc-119(ed3);oxTi880, gcs-1::GFP* (*ldIs3*) and *gst-4::GFP (dvIs19)* were obtained from Caenorhabditis Genetics Center (CGC) (www.cgc.umn.edu). Strain maintenance and experiments were performed at 20°C.

The GFP::TAP plasmids were co-injected with pVB641OB (each at 50 ng/uL) to generate strains to assay TAP localization to the ER. *svIs135* expressed GFP::SEC-61 β, *svEx917* expressed GFP::CYTB-5.1, *rawEx14* expressed GFP::SERP-1.1 and *rawEx21* expressed GFP::SEC-61β^ΔTMD^.

The *asna-1(ΔHis164)* animals were created by deletion of Histidine in position 164 using Crispr-CAS9 technology by Suny Biotech and was maintained over the balancer *hT2(qIs48)*. The ASNA-1^ΔHis164^::GFP expressing transgene, *svEx591* has been described previously (Hemmingsson et al., 2010).

### TAP-targeting analysis

Live animals were sedated in 1 mM Levamisole/M9 and mounted onto 2 % agarose pads. The fluorescence signals of the samples were subsequently analysed at 488 nm and 555 nm by the Confocal Laser Scanning Microscope (LSM700, Carl Zeiss) with LD C-Apochromat 40x/1.1 W Corr. objective. Image processing of Z-stacks was performed in the ZEN Lite (Carl Zeiss) program while the correlation quantification was done in the image analysis software Volocity (PerkinElmer). Correlation quantification was done using Automatic Thresholding (Costes et al., 2004) method to set thresholds objectively.

### Membrane preparations

Synchronized animals were grown to young adults in standard conditions at 20°C. Harvested animals were washed two times in M9 buffer and lysed in extraction buffer (50 mMTris, pH7.2, 250 mM sucrose, 2 mM EDTA) with a bullet blender. Carcasses were immediately spun down at 2900 × g at 4 °C. Supernatants were then collected and spun for 60 min at 100 000 × g at 4 °C. The supernatant fraction was concentrated using Vivaspin Concentrators (Sigma). The pellet fraction was resuspended in RIPA buffer. Pellet and supernatant fractions were run on a Tris-glycine gel, blotted onto a polyvinylidene diflouride membrane and probed with the anti-GFP antibody [3H9] (Chromotek) or anti-RME-1 (Hadwiger et al., 2010).

### Insulin assays

Larval arrest phenotypes were scored in the F1 generation, grown at 20°C, from *wrb-1* dsRNA-injected mothers. Worms harboring integrated *daf-16::gfp* and *daf-28::gfp* arrays were grown at 20°C and imaged using a Nikon Ni-E microscope, equipped with Hammamatsu Orca flash4.0 camera. *daf-16::gfp* animals were analysed within 10 minutes to avoid artifacts due to stressful conditions. DAF-28::GFP uptake by coelomocytes was scored in adult *svIs69; ok938*, in *svIs69; wrb-1(RNAi)* worms and in *svIs69;asna-1(A63V)*.

### Western blot analysis

*Reducing SDS-PAGE:* Nematodes were synchronized in young adult stage and homogenized in a T-PER Tissue Protein Extraction Reagent (Thermo Scientific) with Halt Protease and Phosphatase Inhibitor Cocktail (Thermo Scientific). Homogenization was done using the Next Advance cell disruptor and 0.2mm stainless steel beads for 3 minutes at 4°C. Bradford protein assay was performed to measure protein concentration. Samples were boiled for 10 min in reducing loading buffer (SDS and β-mercaptoethanol). Mini-PROTEAN TGX stain free gradient precast gels (BioRad) were used for gel electrophoresis in Tris/Glycine/SDS Buffer (BioRad) and Trans-Blot Tubro Transfer System Transfer pack membranes (BioRad) for transfer to PVDF membranes. *Non-reducing SDS-PAGE:* Protein samples isolated form the nematodes were boiled for 10 min in non-reducing (without β-mercaptoethanol) loading buffer prior SDS-PAGE analysis. SDS-PAGE analysis was performed as described above. *Antibodies:* anti-ASNA-1 antibody, 1:1000 and anti-GFP antibody [3H9] Chromotek, 1:1000. To confirm equal loading, membranes were stripped using Restore Plus (Thermo Scientific) and incubated with anti-alpha tubulin [T5168] (Sigma). Chemiluminescent signals were generated using the Supersignal West Femto detection reagent (Thermo Scientific) and detected using the LAS1000 machine equipped with a cooled CCD camera (Fuji). Each panel in western blot figures is composed of lanes from the same individual gel in which extra lanes between samples were removed.

### Pro-oxidant, anti-oxidant treatment and cisplatin treatment

*H_2_O_2_/CuSO_4_*: Adult worms were washed from the plate 3 times with M9 and incubated in dark with pro-oxidant: 10 mM H_2_O_2_ and 1mM CuSO_4_ simultaneously for 30 min. Worms were washed 3 times with M9 followed by protein isolations. *Sodium arsenite* (Sigma): Adult worms were washed 3 times with M9 and transferred into the solution containing 5 mM sodium arsenite and incubated for 1h. Worms were washed 3 times with M9 followed by protein isolation. *MitoTempo* (Sigma): Adult worms were staged by gravity and L1 larvae were collected. Larvae were grown on plates with OP50 for 24h and transferred into plates containing 50 µM mitoTempo Worms grow on plates with mitoTempo seeded with OP50 bacteria for 48h. Worms were washed 3 times with M9 followed by protein isolation. *Epigallocatechin gallate (EGCG)*: EGCG (Sigma) containing plates were prepared by spreading 400 µM EGCG on the unseeded NGM plate to obtain a final concentration of 28.6 µM. Spots of concentrated OP50 was used as a source of food after the plates had dried. Mixed stage worms washed and synchronized. The supernatant containing L1 larvae was collected. Larvae were grown on plates with OP50 for 24h and L4 larvae from these plates were transferred onto EGCG plates. Worms grown on plates with EGCG and OP50 for 24h and were transferred onto a fresh EGCG plate for another 24h. Worms were washed 3 times with M9 followed by protein isolation. *Cisplatin treatment:* Cisplatin plates were prepared using MYOB media with 2% agar in which the drug was added at a concentration of 300μg/mL. Cisplatin solution (Accord Healthcare AB) was added to autoclaved medium after cooling down to 52°C. Adult worms were washed from the plate 3 times with M9 and incubated on cisplatin for 3h. Worms were washed from the cisplatin plates for the protein insolation.

### Transmission electron microscopy (TEM)

Worms were washed 3x in M9. The anterior portion of the head was cut off in fixative solution, 2.5 % (v/v) glutaraldehyde in 0.1 M cacodylate. Samples were then incubated overnight in fixative at 4 °C. After washing three times in fixative solution, samples were treated with 1 % (v/v) osmium tetroxide for 1 h and then washed twice in distilled water. Dehydration with 50, 70, 95, and 100 % ethanol was followed by infiltration and embedment in Spurr’s resin. Using a DiATOME diamond knife on a Leica EM UC7, sections (60-90 nm) were mounted on copper grids. The grids were then treated with 5 % uranyl acetate in water for 20 minutes, followed by Sato’s lead staining for 5 minutes. Sections were examined with a Jeol 1230 transmission electron microscope and images were captured using a Gatan MSC 600CW camera.

### Cisplatin sensitivity assay

Cisplatin plates were prepared using MYOB media with 2 % agar in which the drug was added at a concentration of 300 μg/mL or 500 μg/mL. Cisplatin solution from a stock at 1 mg/mL (Accord Healthcare AB) was added to autoclaved medium after cooling it to 52°C. L4 larvae were picked to a new plate and allowed to grow for 24h before exposure to cisplatin. Cisplatin exposure was for 24 ± 1h at 20°C and death was determined by absence of touch-provoked movement when worms were stimulated by a platinum wire.

### RNA isolation and quantitative PCR

Synchronized young adult worms were washed with M9 solution and transferred into RNase free tubes. Total RNA was extracted from worms suspended in around 25 μL of M9 and 360 μL of NucleoZOL (Macherry-Nagle) and disrupted by six freeze-thaw cycles between a dry ice/ethanol bath and a 37°C water bath. Aurum Total RNA Mini Kit (BioRad) was used; cDNA was synthesized using iScript cDNA Synthesis Kit (BioRad). qPCR was performed on a StepOnePlus Real-Time PCR System (Applied Biosystems) instrument using KAPA SYBR FAST qPCR Kit (KapaBiosystems) with the comparative Ct method and normalization to the housekeeping gene *F44B9.5*. All samples were tested in triplicate.

### Statistical analysis

Statistical analysis was performed with Prism 7 software (GraphPad software, La Jolla, CA, USA). Statistical significance was determined using a Student’s t-test or One-way ANOVA. P-values <0.05 indicated statistical significance.

### Immunoprecipitation assay

Nematodes were grown on NGM plates with OP50 for 4 days. Worms were washed 3 times with M9, lysed using Next Advance cell disruptor with 0.2mm stainless steel beads for 3 minutes at 4°C in lysis buffer (10mM Tris/Cl, 150mM NaCl, 0.5mM EDTA, 0,5% NP-40). Bradford protein assay was performed to measure protein concentration. Lysate was added to 35µl of GFP-Trap®_MA magnetic beads (Chromotek) and tumbled end-over-end for 1 hour at 4°C. Beads were magnetically separated and washed 3 times with 500µl of wash buffer I (0,01% Tween-20 in 50mM TEAB) for 10 min, followed by 3 times wash II (50mM TEAB) for 10 min. Elution of protein from the beads was performed by adding 50 µl 0.2M glycine (pH 2.5) for 30 sec, followed by magnetic separation. 5 µl 1M Tris base (pH10.4) was added for neutralization. GFP expressing strain (*unc-119(ed3);oxTi880*) was used as a negative control for GFP-interacting partners. Experiment was performed three times for identification of WRB-1 as an interacting partner.

### Carbonylated protein detection

Protein carbonylation was determined by OxyBlot Protein Oxidation Detection Kit (Millipore). Wild-type animals were synchronized to the young adult stage. Worms were exposed to cisplatin at concentration of 300 μg/mL for 3 or 6 hours. Next worms were washed off the plate using M9 solution. Worms were homogenized as described above. Protein estimation was done using Bradford assay (Bio-Rad). Total extracted proteins (20 μg) were derivatized according to the manufacturer’s instructions (Millipore). 2-mercaptoethanol was used to reduce samples. The membrane was blocked for 1 hour in 1% BSA/TBS-T and probed with antibodies provided in the kit according to the manufacturer’s instructions (Millipore).

### Estimation of oxidative stress

Reactive oxygen species (ROS) formation was quantified using 2,7-dichlorodihydrofluorescein diacetate (H_2_DCFDA) in the control vs. cisplatin treated worms. *For cisplatin treated animals*: Approximately 1000 worms were transferred to the cisplatin plate and incubated for 3h (with concentrated OP50 as a source of food). After the incubation worms were washed off and cleaned from the bacteria and left in about 100 μL M9, then transferred to assay well of 96 well plate. 100 μL of 100 μM of 2,7-dichlorodihydrofluorescein diacetate (H_2_DCFDA) (Thermo Fisher Scientific) was added to achieve a final concentration of 50 μM H_2_DCFDA. *For control animals*: Approximately 1000 worms in about 100 μL M9 were transferred to assay well of 96 well plate and addition of 100 μL of 100 μM 2,7-dichlorodihydrofluorescein diacetate (H_2_DCFDA) (Thermo Fisher Scientific), so as to achieve a final concentration of 50 μM H_2_DCFDA. Fluorescence was read at the time of adding the dye and one hour after addition of dye, using fluorimeter {485 excitations, 520 emissions}. Initial readings were subtracted from the final readings and fluorescence per 1000 worms was calculated.

## ACKNOWLEDGEMENTS

We thank the Caenorhabditis Genetic Center (funded by NIH Office of Research Infrastructure Programs P40 OD010440) and National Bioresource Project for the Experimental Animal “Nematode C. elegans” for providing strains, the Centre for Cellular Imaging at the University of Gothenburg and the National Microscopy Infrastructure, NMI (VR-RFI 2016-00968) for providing assistance in microscopy, Proteomics Core Facility of Sahlgrenska Academy, University of Gothenburg for proteomic analysis, members of Simon Tuck and Marc Pilon labs for scientific discussions and G. Wolfstetter, K. Pfeifer, J. Nilsson and L. Nilsson for helpful comments on the manuscript.

## Author contributions

D.R., O.B., A.P., G.K., O.H. and P.N. designed experiments. D.R., O.B., A.P. and G.K. performed the experiments. D.R., O.B., A.P. and G.K. analyzed the data. D.R., O.B., A.P., G.K., O.H. and P.N. wrote the paper. All authors discussed the results and conclusions in the manuscript

## DATA AND MATERIALS AVAILABILITY

All data is available in the main text or the supplementary materials.

## SUPPLEMENTARY FIGURES

**Figure S1.**
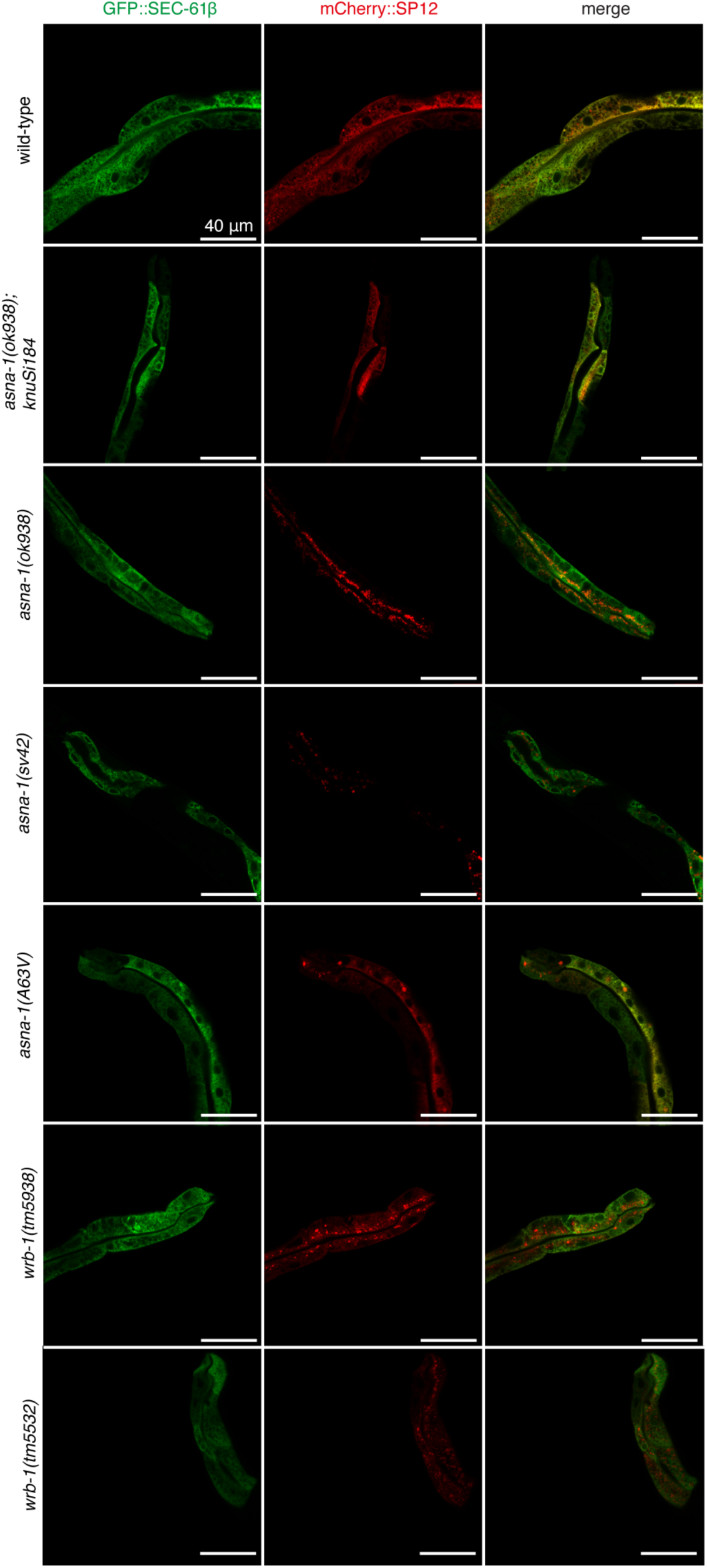
ASNA-1 and WRB-1 promote ER targeting of TAPs *in vivo*. Confocal imaging of wild-type, *asna-1(ok938);knuSi184*, *asna-1(ok938)*, *asna-1(sv42)*, *asna-1(A63V), wrb-1(tm5938)*, or *wrb-1(tm5532)* and animals co-expressing GFP::SEC-61β and mCherry::SP12. In all cases, posterior intestinal cells were imaged.

**Figure S2.**
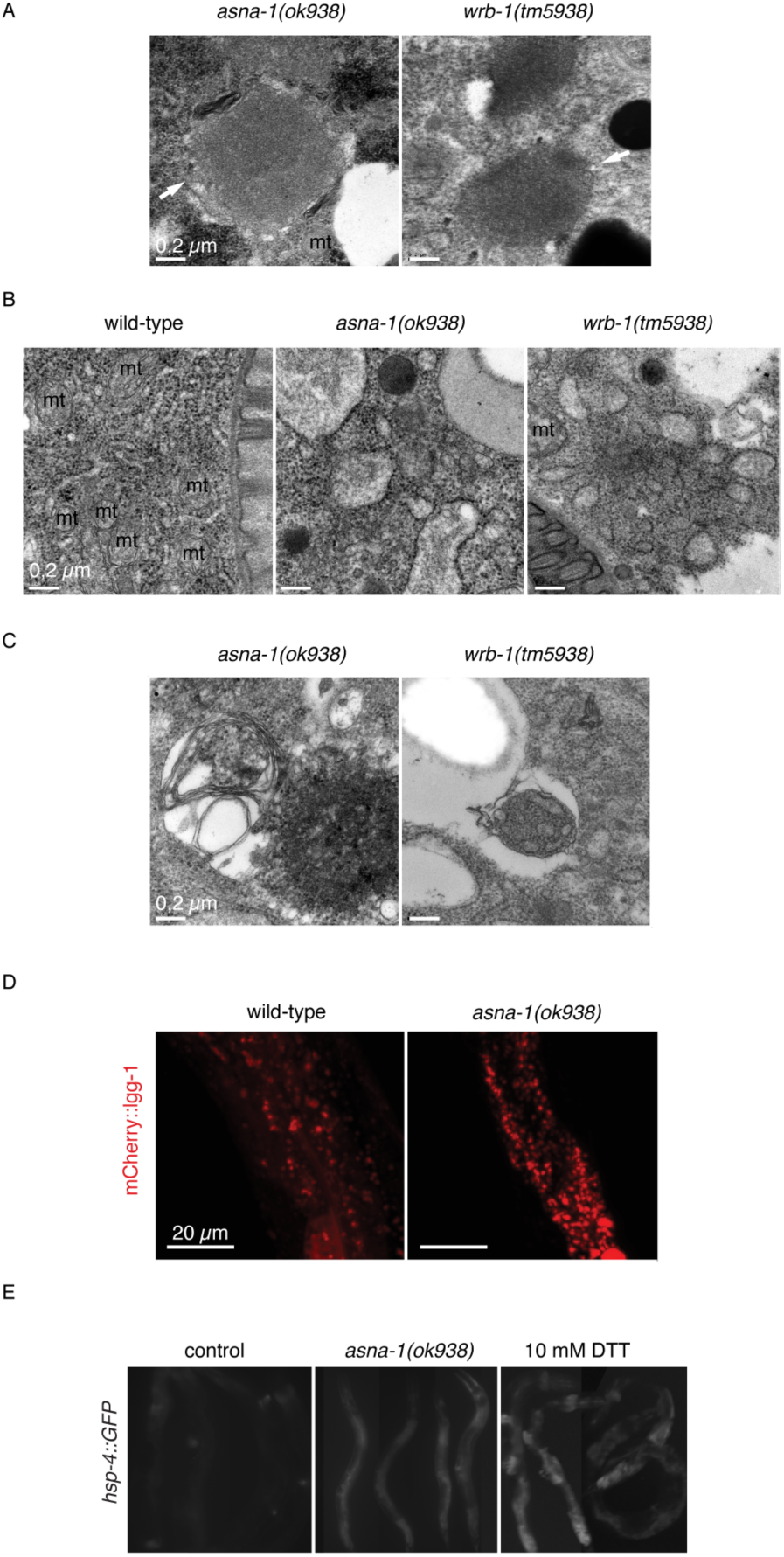
Ultrastructural phenotypes of *asna-1(−) and wrb-1(−)* animals and autophagy and ER stress levels in *asna-1* mutants. (A) Examples of numerous cytosolic inclusion bodies (white arrows) found in the intestinal cells of *asna-1(ok938)* and *wrb-1(tm5938)* animals. The black arrowheads highlight examples of membrane whorls that in some cases flank the inclusion bodies in intestinal cells of *asna-1(ok938)* animals. (B) Dilated ER lumina in intestinal cells of *asna-1(ok938)* and *wrb-1(tm5938)* animals. Black arrowheads indicate rough ER (RER) membranes. (C) Portions of RER were found to be engulfed in ER-containing autophagosomes (ERAs) in *asna-1(ok938)* and *wrb-1(tm5938)* animals. D) Confocal images of representative adult animals co-expressing mCherry::lgg-1. LGG-1 localization in wild-type animals (left panel) is more diffuse and less punctate compared to *asna-1(ok938)* animals (right panel). (E) Expression from the Phsp-4::GFP reporter imaged by fluorescence microscopy. The reporter induction upon introduction of *asna-1(ok938)* mutation is comparable to the 10 mM DTT exposure for 4 h. Animals undergoing the L3/L4 molt were excluded from the analysis, since molting caused a sharp increase in Phsp-4::GFP expression.

**Figure S3.**
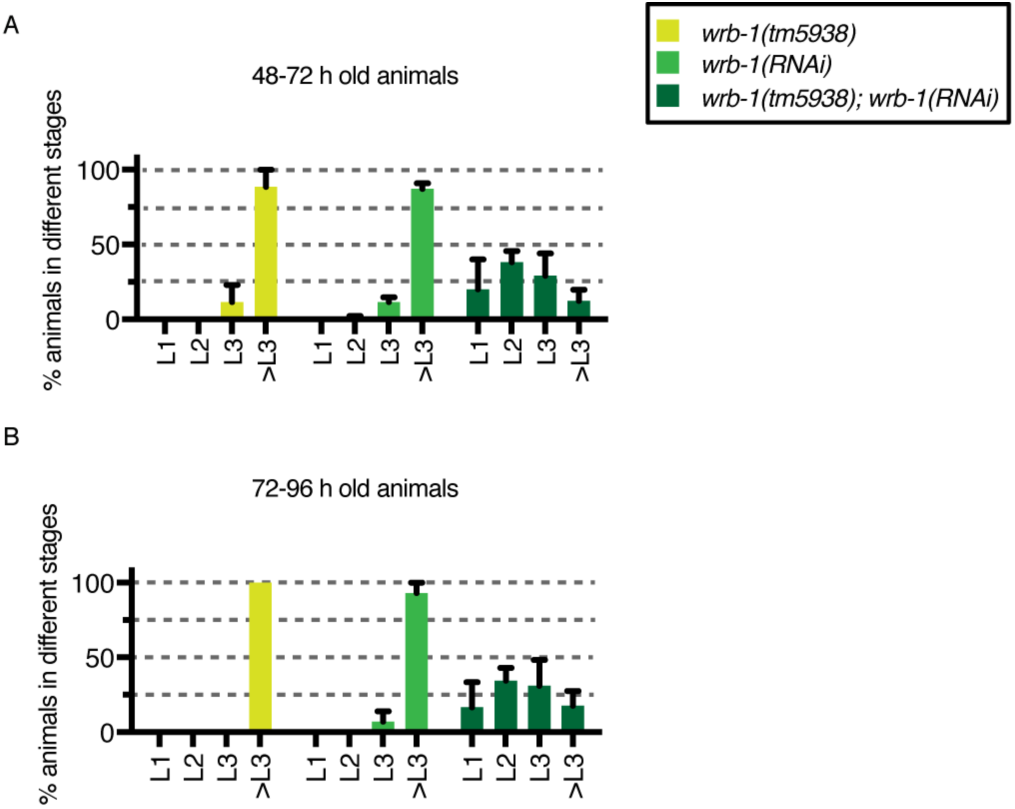
Depleting both maternal and zygotic *wrb-1*(*m^−^z^−^)* did not reproduce the reversible arrest in the first larval stage characteristic for *asna-1*(*m^−^z^−^)* *wrb-1(tm5938)* animals were injected with *wrb-1* dsRNA into the gonads and allowed to lay eggs for 24 h before being removed. Animals were scored after 48 h (A) and 72 h (B) by counting the number of animals in each stage. Bars represent mean ± SD.

**Figure S4.**
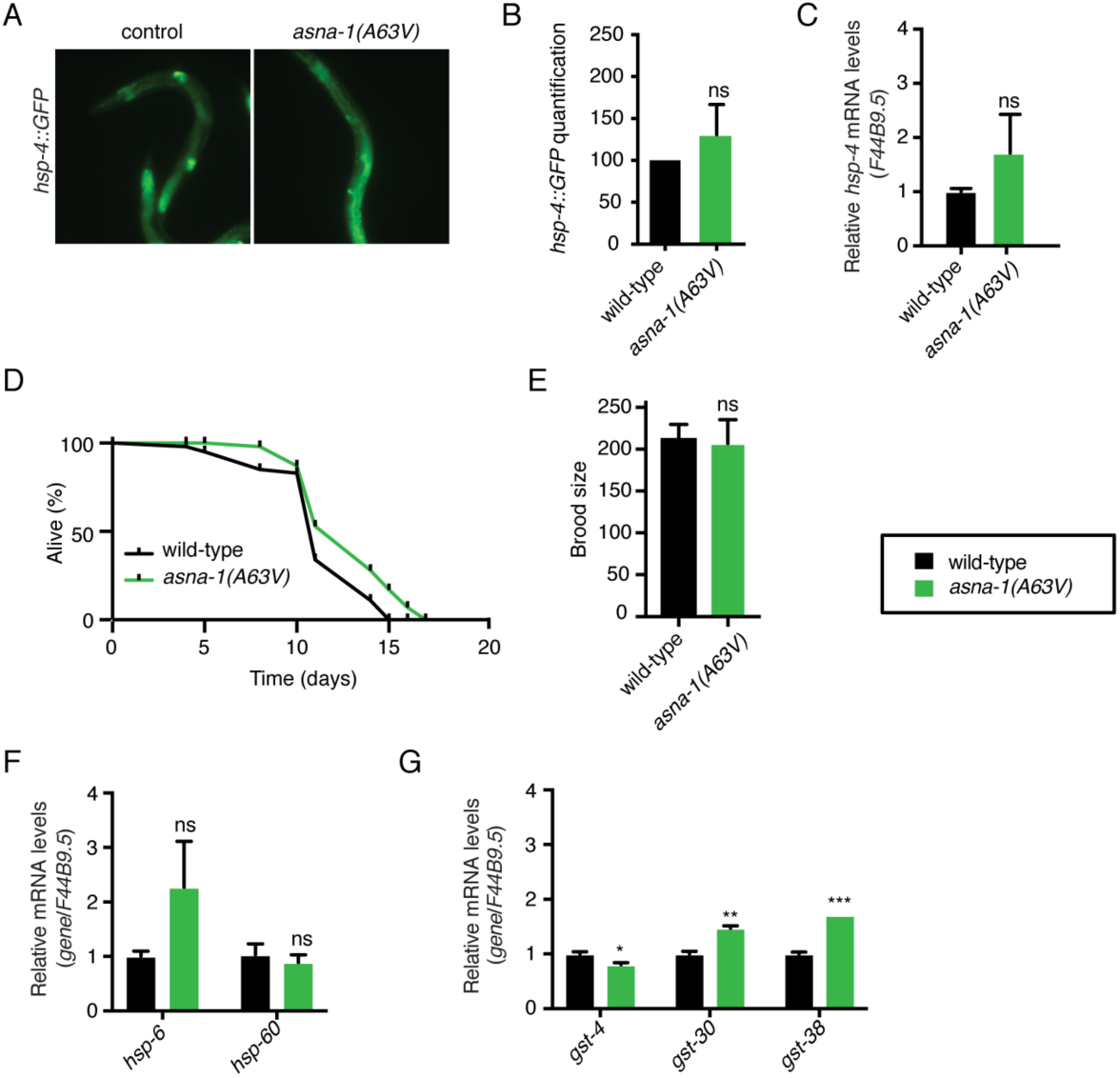
Cellular stress response analysis in *asna-1(A63V)* (A) Expression from the Phsp-4::GFP reporter imaged by fluorescence microscopy in the wild-type and *asna-1(A63V)* animals. (B) Phsp-4::GFP expression quantification in the wild-type and *asna-1(A63V)* animals (n≥6). Statistical significance was determined by the independent two-sample t-test (*p<0.05, **p<0.01, ***p<0.001). Bars represent mean ± SD. (C) Relative mRNA analysis of ER stress reporter *hsp-4* in *asna-1(A63V)* animals. Statistical significance was determined by the independent two-sample t-test (*p<0.05, **p<0.01, ***p<0.001). Experiments were performed in triplicate (n=3). F44B9.5 was used as a normalizing control. Bars represent mean ± SEM. (D) Life span analysis of wild-type and *asna-1(A63V)* animals (n≥42). (E) Brood size analysis of wild-type and *asna-1(A63V)* (n=5). Statistical significance was determined by the independent two-sample t-test (*p<0.05, **p<0.01, ***p<0.001). Bars represent mean ± SD. (F) Relative mRNA analysis of the mitochondrial stress reporters (*hsp-6* and *hsp-60*) and (G) oxidative stress reporters (*gst-4*, *gst-30* and *gst-38*) in *asna-1(A63V)* animals. Statistical significance was determined by the independent two-sample t-test (*p<0.05, **p<0.01, ***p<0.001). Experiments were performed in triplicate (n=3). F44B9.5 was used as a normalizing control. Bars represent mean ± SEM.

